# Chronic activation of tubulin tyrosination in HCM mice and in human iPSC-engineered heart tissues improves heart function

**DOI:** 10.1101/2023.05.25.542365

**Authors:** Niels Pietsch, Christina Y. Chen, Svenja Kupsch, Lucas Bacmeister, Birgit Geertz, Marisol Herrera-Rivero, Hanna Voß, Elisabeth Krämer, Ingke Braren, Dirk Westermann, Hartmut Schlüter, Giulia Mearini, Saskia Schlossarek, Jolanda van der Velden, Matthew Caporizzo, Diana Lindner, Benjamin L. Prosser, Lucie Carrier

**Affiliations:** Department of Experimental Pharmacology and Toxicology, University Medical Center Hamburg-Eppendorf, Hamburg, Germany; DZHK (German Centre for Cardiovascular Research), partner site Hamburg/Kiel/Lübeck, Hamburg, Germany; Department of Physiology, Pennsylvania Muscle Institute, University of Pennsylvania Perelman School of Medicine, Philadelphia, PA 19104, USA; Gene Therapy Program, Department of Medicine, Perelman School of Medicine, University of Pennsylvania, Philadelphia, PA, USA (current affiliation); Department of Cardiology, University Heart and Vascular Center, University Medical Center Hamburg-Eppendorf, Hamburg, Germany; Institute of Experimental Cardiovascular Research, University Medical Center Hamburg-Eppendorf, Hamburg, Germany (current affiliation); Department of Cardiology and Angiology, University Heart Center Freiburg-Bad Krozingen, Faculty of Medicine, University of Freiburg, Freiburg, Germany (current affiliation); Department of Genetic Epidemiology, Institute of Human Genetics, University of Münster, Münster, Germany; Department of Psychiatry, University of Münster, Münster, Germany (current affiliation); Section Mass Spectrometric Proteomics, University Medical Center Hamburg-Eppendorf, Hamburg, Germany; Vector facility, Department of Experimental Pharmacology and Toxicology, University Medical Center Hamburg-Eppendorf, Hamburg, Germany; DiNAQOR AG, 8952 Schlieren, Switzerland (current affiliation); Amsterdam UMC, Vrije Universiteit Amsterdam, Department of Physiology, Amsterdam Cardiovascular Sciences, Amsterdam, The Netherlands; Department of Molecular Physiology and Biophysics, University of Vermont Larner College of Medicine, Burlington, VT, USA (current affiliation)

**Keywords:** heart function, hypertrophic cardiomyopathy, engineered heart tissue, hiPSC, tubulin tyrosination and detyrosination

## Abstract

**Rationale:** Hypertrophic cardiomyopathy (HCM) is the most common cardiac genetic disorder caused by sarcomeric gene variants and associated with left ventricular (LV) hypertrophy and diastolic dysfunction. The role of the microtubule network has recently gained interest with the findings that α-tubulin detyrosination (dTyr-tub) is markedly elevated in heart failure. Acute reduction of dTyr-tub by inhibition of the detyrosinase (VASH/SVBP complex) or activation of the tyrosinase (tubulin tyrosine ligase, TTL) markedly improved contractility and reduced stiffness in human failing cardiomyocytes, and thus poses a new perspective for HCM treatment.

**Objective:** In this study, we tested the impact of chronic tubulin tyrosination in a HCM mouse model (*Mybpc3*-knock-in; KI), in human HCM cardiomyocytes and in SVBP-deficient human engineered heart tissues (EHTs).

**Methods and Results:** AAV9-mediated TTL transfer was applied in neonatal wild-type (WT) rodents and 3-week-old KI mice and in HCM human induced pluripotent stem cell (hiPSC)-derived cardiomyocytes. We show that i) TTL for 6 weeks dose-dependently reduced dTyr-tub and improved contractility without affecting cytosolic calcium transients in WT cardiomyocytes; ii) TTL for 12 weeks improved diastolic filling, cardiac output and stroke volume and reduced stiffness in KI mice; iii) TTL for 10 days normalized cell hypertrophy in HCM hiPSC-cardiomyocytes; iv) TTL induced a marked transcription and translation of several tubulins and modulated mRNA or protein levels of components of mitochondria, Z-disc, ribosome, intercalated disc, lysosome and cytoskeleton in KI mice; v) SVBP-deficient EHTs exhibited reduced dTyr-tub levels, higher force and faster relaxation than TTL-deficient and WT EHTs. RNA-seq and mass spectrometry analysis revealed distinct enrichment of cardiomyocyte components and pathways in SVBP-KO vs. TTL-KO EHTs.

**Conclusion:** This study provides the first proof-of-concept that chronic activation of tubulin tyrosination in HCM mice and in human EHTs improves heart function and holds promise for targeting the non-sarcomeric cytoskeleton in heart disease.

## Introduction

Hypertrophic cardiomyopathy (HCM) is the most common inherited cardiac disease caused by mutations in several sarcomeric genes. *MYBPC3*, encoding cardiac myosin-binding protein C is the most frequently mutated gene.^1^ HCM is associated with left ventricular (LV) hypertrophy, diastolic dysfunction and increased interstitial fibrosis. Treatment of severe HCM with outflow tract obstruction is limited to surgical removal of septal tissue or catheter ablation.

The role of the microtubule network during progression of cardiac diseases has gained interest in recent years with the findings that microtubule detyrosination and associated cardiomyocyte stiffness is elevated in HCM and heart failure^2, 3^ (for reviews,^4-7^). The cycle of α-tubulin detyrosination/re-tyrosination consists in the catalytic removal and re-incorporation of the last C-terminal tyrosine of α-tubulin. We have previously reported higher levels of detyrosinated tubulin (dTyr-tub) in human and mouse models of HCM.^3, 8^ The tubulin tyrosine ligase (TTL) catalyses the tyrosination, and the tubulin carboxypeptidase complex, composed of vasohibin (VASH1 or VASH2) and its chaperone, the small vasohibin-binding protein (SVBP), catalyses the removal of the C-terminal tyrosine. VASH1 is the major isoform in human heart.^9^ Recently, it was shown that an acute reduction of dTyr-tub by TTL overexpression or *VASH1* knockdown markedly improved contractility and reduced stiffness in human failing cardiomyocytes^8, 9^ and thus poses a possible new perspective for the treatment of HCM.

In this study, we evaluated the impact of long-term adeno-associated virus serotype 9 (AAV9)-mediated TTL overexpression in neonatal wild-type (WT) rodents and in 3-week-old homozygous *Mybpc3*-targeted knock-in (KI) mice that present with eccentric LV hypertrophy and both systolic and diastolic dysfunction.^10, 11^ Using a combination of approaches, we show that a long-term TTL overexpression lowered dTyr-tub, improved contractility without affecting calcium transients in WT rat cardiomyocytes, and improved diastolic filling, compliance, stroke volume and cardiac output in KI mice. We further provide evidence that TTL overexpression did not significantly affect the fetal gene program and LV hypertrophy, whereas it modulated mRNA or protein levels of components of mitochondria, Z-disc, ribosome, intercalated disc, lysosome and cytoskeleton in KI mice. These data were supported by a normalization of cell hypertrophy after a 10-day TTL gene transfer in HCM induced pluripotent stem cell (hiPSC)-derived cardiomyocytes, and by SVBP deficiency in hiPSC-derived cardiomyocytes and engineered heart tissues (EHTs).

## Methods

### Animals

The investigation conforms to the guidelines for the care and use of laboratory animals published by the NIH (Publication No. 85-23, revised 1985). For studies in WT C57BL/6J mice and Sprague-Dawley rats, animal care and use procedures were approved and performed in accordance with the standards set forth by the University of Pennsylvania Institutional Animal Care and Use Committee and the Guide for the Care and Use of Laboratory Animals published by the US National Institutes of Health. KI mice were generated previously,^10^ and both KI and WT mice were maintained on a Black Swiss background. The experimental procedures performed in KI and WT mice were in accordance with the German Law for the Protection of Animals and accepted by the Ministry of Science and Public Health of the City State of Hamburg, Germany (Nr. 074-19).

### Human samples

The HCM patient carrying the *MYBPC3* (c.2308G>A; p.Asp770Serfs98X) mutation was recruited in the outpatient HCM clinic at the University Heart and Vascular Center Hamburg and provided written informed consent for genetic analysis and the use of skin fibroblasts ^12^. All procedures were in accordance with the Code of Ethics of the World Medical Association (Declaration of Helsinki). The study was reviewed and approved by the Ethical Committee of the Ärztekammer Hamburg (PV3501).

### Plasmid constructs and AAV9 particle production

The TTL-IRES-dsRed insert was cloned into an expression construct flanked by AAV2 inverted terminal repeats containing a human cardiac troponin T (TNNT2) promotor, a chimeric intron, a post-transcriptional regulatory element (WPRE) and a rabbit beta-globin poly A sequence (PENN.AAV.cTNT.TTL.IRES.dsRed; Figure S1A). AAV9 pseudotyped vector was generated by triple transfection of HEK293 cells and iodixanol purification (Penn Vector Core, University of Pennsylvania) as previously described.^13^ Genome copy titer was determined using a droplet digital PCR-based method.^14^

The pGG2-*TNNT2*-*HA-TTL* and the pAAV-*TNNT2*-Empty plasmids were generated from the pGG2-*TNNT2*-Mut1 plasmid (VC206) by InFusion Cloning (Takara Clontech). In brief, the full-length HA-TTL was amplified from the HA-TTL-IRES-dsRed plasmid using PrimeStar GLX Polymerase (Takara Clontech) and ligated into the NheI and Bam*H*I restriction sites of VC206. The Empty plasmid was generated by amplification of the *TNNT2* promoter region from VC206 plasmid and inserted into MluI and HindIII restriction sites of pAAV-GFP (Cellbiolabs). After ligation, ampicillin-resistant clones were tested by enzymatic digestion for presence of the insert and one positive clone of each pGG2-*TNNT2*-*HA-TTL* and pAAV-*TNNT2*-Empty was sequenced. AAV9 pseudotyped vectors were generated by triple transfection of HEK293 cells with pGG2-*TNNT2*-*HA-TTL* or pAAV-*TNNT2*-Empty, pE8/9 (kindly provided by Julie Johnston, Penn Vector Core, University of Pennsylvania, PA, USA) and pHelper encoding adenoviral helper functions (Cellbiolabs). Generation of recombinant AAV9 particles was carried out as described previously.^15, 16^

### In vivo AAV9 administration

Neonatal WT C57BL/6 mice received (subcutaneous injection) AAV9-TTL-dsRed at a dose of 1.75E11 or 9E11 vg/g at postnatal day 2 (P2). Neonatal Sprague Dawley rats received AAV9-TTL-dsRed at a dose of 4.4E11 vg/g at P4 (pericardial injection). Three-week-old WT and KI mice received AAV9 encoding either an empty cassette (AAV9-Empty) or HA-tagged TTL (AAV9-TTL) at a dose of 2.25E11 vg/g via systemic administration into the tail vein using a 30-G needle.^15-17^

### Isolation of adult mouse and rat ventricular myocytes and functional assessment

Primary ventricular cardiomyocytes were isolated as previously described.^18^ Briefly, the heart was removed from 6-week-old mouse or rat anesthetized under isoflurane and retrograde-perfused on a Langendorff apparatus with a collagenase solution. The digested heart was then minced and triturated using a glass pipette. The resulting supernatant was separated and centrifuged at 300 revolution per minute (rpm) to isolate cardiomyocytes which were then resuspended in cardiomyocyte media for functional assessment. Elastic modulus was measured using nanoindentation (Piuma Chiaro, Optics11, The Netherlands) as previously described.^19^ Cardiomyocytes contractility and calcium transients were measured as previously described.^9^

### Human iPSC-derived cardiomyocytes and EHT contractility measurements

A control hiPSC line carrying a monoallelic mTag-RFP-T-TUBA1B (clone AICS-0031-035, Allen Institute for Cell Science, Seattle, WA, USA) was expanded and used to create SVBP-KO and TTL-KO hiPSC lines with CRISPR/Cas9 technology. Two sgRNAs were selected for each locus to create a large deletion (Figure S2A). Cells were nucleofected using the Amaxa 4D nucleofector (Lonza) with pre-annealed ribonucleoprotein complexes of sgRNAs and recombinant Cas9 protein (IDT, 1081058), single cell clones expanded and genotyped for successful homozygous deletions (Figure S2B). Clones with large homozygous deletions for each of the genes of interest underwent quality control in karyotyping (Figure S2C) by nCounter profiling (NanoString, Seattle, US), pluripotency validation by SSEA-3 (stage-specific embryonic antigen 3) staining (BD Biosciences, 560892; Figure S2D) and control for mycoplasma contamination. To ensure specific activity of the CRISPR-Cas9 system, the 10 most likely off-target loci were identified with crispor.tefor.net,^20^ and sequenced to assess genome integrity. No alterations were detected, indicating no unspecific activity of the CRISPR-Cas9 complex (data not shown). After expansion of the 3 hiPSC lines, cardiomyocytes were produced according to the monolayer protocol.^21^ RT-qPCR was performed with RNA isolated from hiPSC-CMs to ensure mRNA deficiency of the respective gene (Figure S2E,F). EHTs were cast from 2-6 independent batches of cardiomyocyte differentiation (TNNT2-positive cardiomyocytes > 85%) of the 3 hiPSC lines on standard silicone posts (0.28 mN/mm) or on stiffer silicone that induced afterload enhancement (0.8 mN/mm). Auxotonic contractions were measured until day 60 as previously described.^22-24^

We also used a patient-derived heterozygous *MYBPC3* (MYBPC3het; UKEi070-A) and its isogenic control (MYBPC3ic; UKEi070-A-1) hiPSC lines that were previously created.^12^ MYBPC3het and MYBPC3ic hiPSC-cardiomyocytes (3-4 batches of differentiation of triplicates) were transduced with AAV9-TTL or AAV9-Empty (MOI 100,000) for 10 days and treated with 100 nM endothelin-1 (ET1) or vehicle (H_2_O) the last 3 days as previously described.^25^ Cardiomyocytes were then stained for α-actinin (Sigma, SAB2108642, 1:800) and N-cadherin (Sigma, C2542, 1:500), followed by the secondary antibodies Alexa Fluor^TM^ 488 (mouse) and Alexa Fluor^TM^ 546 (rabbit) (Invitrogen, A11001; A11035, 1:800), and cell area was analysed by confocal microscopy and ImageJ.

### Echocardiographic analysis in mice

Transthoracic echocardiography was performed using the Vevo 3100 System (VisualSonics, Toronto, Canada) as described previously.^15-17^

### Hemodynamic measurements in mice

Hemodynamic measurements were performed using a PV loop system (ADV500, Transonic) in closed-chest approach.^26, 27^ Mice were anesthetized using urethane (0.8-1.2 g/kg body weight) and buprenorphine analgesia (0.1 mg/kg body weight), intubated and artificially ventilated. A pressure-conductance catheter (1.2F, Transonic) was inserted in the right carotid artery and carefully pushed forward into the LV. Catheter’s position inside the ventricle was optimized until rectangular-shaped loops were obtained. PV loops were conducted under short-time apnoea. During the PV loop measurements, inferior vena cava (IVC) was occluded by gentle compression. To estimate the volume, a bolus of hypertonic saline (10%) was injected into the left jugular vein.^28^ Data were acquired using iox2 (Emka Technologies). Heart rate, maximum rate of LV pressure development in systole (dP/d_tmax_), maximal rate of LV pressure development in diastole (dP/dt_min_), LV end-diastolic pressure (LVEDP) and volume (LVEDV), and LV end-systolic pressure (LVESP) and volume (LVESV) were measured. Subsequent analyses of PV loops were performed in LabChart 7.3Pro (AD Instruments). For baseline analysis, 5-10 consecutive loops during end-expiratory ventilation pause were selected to calculate preload-dependent parameters. Preload-independent parameters were analysed by selecting loops during IVC occlusion.^28^ Only mice that underwent appropriate levels of anaesthesia, did not experience excessive bleeding during the procedure, and had technically sufficient measurements were included in the analysis

### Protein extraction and Western blot analysis

For experiments performed in neonatal mice, Western blot of whole heart protein extraction was performed to quantify TTL (1:500; Proteintech, 13618-1-AP), α-tubulin (1:1000; clone DM1A, Cell Signaling, 3873), dTyr-tub (1:1000; Abcam, ab48389) and dsRed protein levels as previously described.^9^ For experiments performed in 3-week-old mice, we used cardiac cytosolic or cytoskeletal-enriched protein fractions as described previously.^15, 16, 29^ For experiments performed in hiPSC-cardiomyocytes or EHTs, we used crude protein extracts. Proteins were separated on 12% SDS-polyacrylamide (29:1) or 4-15% precast polyacrylamide mini-gels (Bio-Rad) and electrotransferred on nitrocellulose membranes. Primary antibodies were directed against the HA epitope (1:2000; Sigma, H3663), dTyr-tub and α-tubulin (1:2,000 and 1:5,000; both kindly provided by Marie-Jo Moutin, Grenoble, France), α-tubulin (1:1,000, Cell Signaling), dTyr-tub (1:1,000, Abcam), α-actinin (ACTN2; 1:20,000; Sigma) or GAPDH (1:10,000; HyTest). Secondary antibodies were peroxidase-conjugated anti-mouse (1:20,000, Dianova; 1:10,000, Sigma), anti-rat (1:10,000 Dianova) or anti-rabbit (1:6,000, Sigma). Proteins were visualized with Clarity Western ECL substrate (Bio-Rad) and signals were detected with the ChemiGenius² Bio Imaging System.

### Titin isoform analysis on mouse cardiac crude protein fractions

Titin isoforms were separated on a 1% w/v agarose gel and stained with SYPRO Ruby protein gel stain (Invitrogen, ThermoFisher Scientific) as previously described.^30^ All samples were measured in triplicate and the average of triplicate measurements per sample was shown.

### Immunofluorescence analysis of hiPSC-cardiomyocytes

2D-culture of cardiomyocytes from the 3 hiPSC lines were seeded onto black, geltrex-coated 96-well plates (ThermoFisher Scientific, 165305) with 90.000 cells/cm^2^ and cultured in low glucose DMEM-based medium for 30 days until cells were washed with PBS and fixed with ROTI®Histofix (Carl Roth, P087.1) for 10 min at 4 °C. For immunofluorescence imaging, cells were incubated with blocking buffer (PBS containing 3 % milk and 0.1 % Triton X-100) and primary antibodies against ACTN2 (1:800, Sigma, A7811) and dTyr-tub (1:1000, kindly provided by Marie-Jo Moutin, Grenoble, France) overnight. Cells were washed with PBS and incubated with secondary antibodies AF-488 rabbit and AF-647 mouse (1:800, ThermoFisher Scientific) in blocking buffer for 90 min at RT. Hoechst 33342 (Invitrogen, H3570) was added 1:2500 for 20 min to stain DNA. Image acquisition was performed with a Zeiss LSM 800 confocal microscope.

### RNA-sequencing

For RNA-sequencing (RNA-seq), total RNA was isolated from LV samples of female WT-Empty, KI-Empty, and KI-TTL mice (N=3 per group) using the Direct-zol RNA Microprep Kit (Zymo Research) followed by a DNase digestion step. Enrichment of mRNAs with the NEBNext Poly(A) mRNA Magnetic Isolation Module was followed by library preparation using the NEBNext Ultra II RNA Directional Library Prep Kit for Illumina (New England BioLabs). Single read sequencing took place on a NextSeq 2000 System (Illumina), using the corresponding NextSeq 2000 P3 Reagents (50 cycles), with a read length of 72 base pairs. The integrity of the RNA and quality of the library were assessed using a TapeStation 4200 (Agilent).

The sequencing data was automatically demultiplexed using the Illumina BCL Convert Software v3.8.2. FastQ files underwent pre-trimming and post-alignment quality control rounds using FastQC v0.11.7 (https://www.bioinformatics.babraham.ac.uk/projects/fastqc/). Removal of Illumina adapters and low-quality sequences was performed with Trimmomatic v0.38.^31^ Reads of length <15 bases, as well as leading and/or trailing bases with quality <3 or no base call, and bases with average quality <15 in a 4-base sliding window were removed. Alignment was performed with HISAT2 v2.1,^32^ using the mouse genome assembly mm10 (Mus musculus, GRCm38). Mapped reads (primary alignments) were sorted by read name using SAMtools v1.8,^33^ and read counts were calculated with HTSeq v0.11.2.^34^

Differential expression was assessed using DESeq2^35^ for all possible comparisons. Raw read counts were filtered to remove genes with less than 10 counts prior to analysis. Here, statistical significance of coefficients was tested using a negative binomial generalized linear model with the Wald test and p-values were adjusted for multiple comparisons following the Benjamini-Hochberg method (Padj). Genes were considered differentially expressed at P or Padj < 0.05 values.

To provide a biological context, each list of differentially expressed genes (DEGs) was subjected to functional enrichment analysis using the web tool g:GOSt from g:Profiler.^36^ All gene ontology (GO) and pathways gene set categories available for Mus musculus or Homo sapiens within this tool were retrieved. Gene sets were considered significantly enriched following a hypergeometric test for overrepresentation within annotated genes, corrected for multiple comparisons using the Benjamini-Hochberg method (Padj<0.05).

### Mass spectrometry

Mouse LV tissue were lysed in 100 mM triethyl ammonium bicarbonate (TEAB) and 1% w/w sodium deoxycholate (SDC) buffer, boiled at 95 °C for 5 min. Samples were sonicated with a probe sonicator for 6 pulses to destroy DNA/RNA. For proteome analysis 20 µg of protein of each sample was taken and disulfide bonds reduced in the presence of 10 mM dithiothreitol (DTT) at 60 °C for 30 min. Cysteine residues were alkylated in presence of 20 mM iodoacetamide at 37 °C in the dark for 30 min and tryptic digestion (sequencing grade, Promega) was performed at a 100:1 protein to enzyme ration at 37 °C overnight. Digestion was stopped and SDC precipitated by the addition of 1% v/v formic acid (FA). Samples were centrifuged at 16,000 g for 5 min and the supernatant was transferred into a new tube. Samples were dried in a vacuum centrifuge.

Samples were resuspended in 0.1% FA and transferred into a full recovery autosampler vial (Waters). Chromatographic separation was achieved on a nano-UPLC system (Dionex Ultimate 3000 UPLC system, Thermo Fisher Scientific) with a two-buffer system (buffer A: 0.1% FA in water, buffer B: 0.1% FA in ACN). Attached to the UPLC was a reversed phase trapping column (Acclaim PepMap 100 C18 trap; 100 µm x 2 cm, 100 Å pore size, 5 µm particle size, Waters) for desalting; a purification followed by a reversed phase capillary column (nanoEase M/Z peptide BEH130 C18 column; 75 µm x 25 cm, 130 Å pore size, 1.7-µm particle size, Waters). Peptides were separated using a 60-min gradient with increasing ACN concentration from 2%-30% ACN. The eluting peptides were analyzed on a quadrupole orbitrap ion trap tribrid mass spectrometer (Fusion, Thermo Fisher Scientific) in data dependent acquisition (DDA) mode. For DDA analysis, the fusion was operated at top speed mode analyzing the most intense ions per precursor scan (2×10^5^ ions, 120,000 Resolution, 120 ms fill time) within 3 s and were analyzed by MS/MS in the ion trap (HCD at 30 normalized collision energy, 1×10^4^ ions, 60-ms fill time) in a range of 400–1300 m/z. A dynamic precursor exclusion of 20 s was used.

### NanoString RNA analysis

Total RNA was extracted from powdered whole heart tissue samples using the SV Total RNA isolation kit (Promega, Madison, WI, USA) according to the manufacturer’s instructions or from hiPSC-cardiomyocyte extracts with the TRIzol^TM^ reagent (Invitrogen, 15596018). RNA concentration, purity and quality were determined using the NanoDrop® ND-1000 spectrophotometer (Thermo Fisher Scientific). For gene expression analysis, customized NanoString’s nCounter® Elements TagSet panels were used (Table S1). About 50 ng RNA of each sample were hybridized to the target-specific capture and reporter probes at 67 °C overnight (16 h) according to manufacturer’s instructions. Samples were cooled down to 4 °C, supplemented with 15 μl H_2_O, and loaded into the NanoString cartridge. Afterwards the nCounter Gene Expression Assay was started immediately. Raw data were analyzed with the nCounter® Sprint Profiler. Transcript levels were determined with the nSolver^TM^ Data Analysis Software including background subtraction using negative controls and normalization to six housekeeping genes (*Abcf1, Actb, Cltc, Gapdh, Pgk1 and Tubb5*). Data are expressed as Log2 ratio or fold-change over the mean of controls, indicated in the legend of the figures.

### Statistical analysis

Data were expressed as mean ± SEM. Statistical analyses were performed by one-way or two-way ANOVA followed by Tukey’s or Dunnett’s post-test, and by unpaired Student’s t-test as indicated in figure legends or text, using the commercial software GraphPad Prism8 (GraphPad Software Inc., Boston, US). A value of P<0.05 was considered statistically significant.

## Results

### Long-term TTL overexpression leads to a dose-dependent reduction in myocyte stiffness and improved contractility in rodent ventricular myocytes

We first evaluated the effect of TTL overexpression on reducing myocardial dTyr-tub level in healthy C57/BL6J mice. For an initial study we tested two different doses of AAV9 (1.75E11 and 9E11 vg/g) encoding TTL-dsRed driven under the *TNNT2* promoter (Figure S1A). Mice were injected at P2 and after 6 weeks of expression myocardial tissue was harvested (Figure 1A). As expected, TTL levels increased in the myocardium in a dose-dependent manner, which was concomitant with decreased dTyr-tub level (Figure 1B). A strong inverse correlation was observed across myocardial samples between increased dsRed and reduced dTyr-tub levels, suggesting that dsRed can be utilized as a useful indicator of dTyr-tub level (Figure 1C). To evaluate transduction efficiency, we isolated cardiomyocytes 6 weeks after a single intermediate dose of AAV9-TTL-dsRed (4.4E11 vg/g) and binned them based on their dsRed fluorescence levels relative to a non-injected control. We found that 85% of myocytes from injected hearts exhibited dsRed levels above that of non-injected controls, with heterogenous levels of expression between transduced myocytes (Figure 1D).

**Figure 1.**
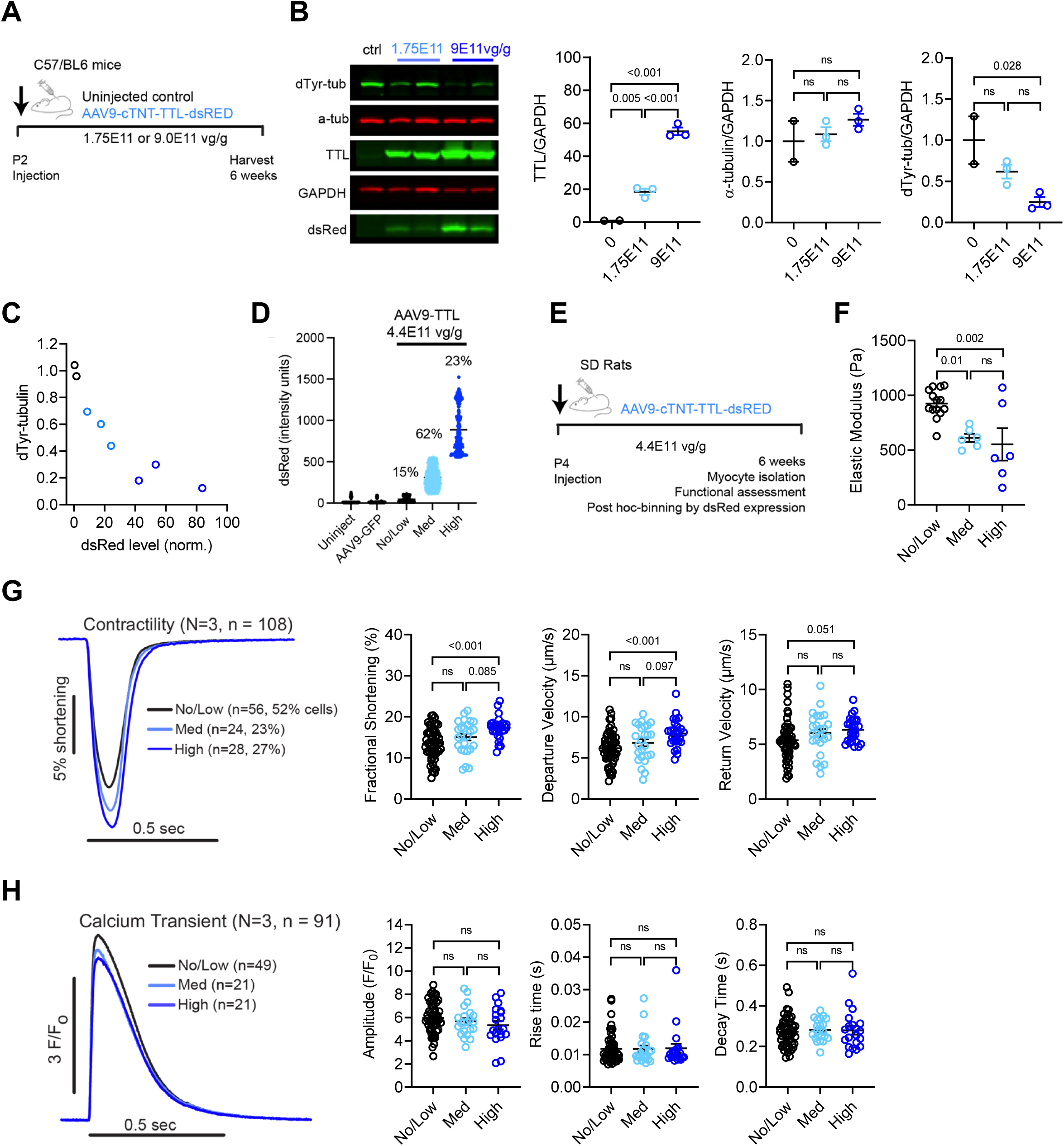
Effect of chronic *TTL* in wild-type rodents and analysis of isolated cardiomyocytes. **A)** Protocol: wild-type neonatal mice (P2) received either an adeno-associated virus serotype 9 (AAV9) encoding TTL-dsRed at a low (1.75E11 vg/g, N=3) or high dose (9E11 vg/g, N=3), or a sham injection (N=2). After 6 weeks, ventricular myocardium was harvested. **B)** Representative Western blot and quantification of TTL, total α-tubulin and detyrosinated tubulin (dTyr-tub), normalized to GAPDH. **C)** Correlation between dsRed protein level (measured via Western blot) and dTyr-tub levels from the same hearts in B. **D)** Percentage of cardiomyocytes expressing no/low, medium or high dsRed intensity after application of AAV9-TTL-dsRed of 4.4E11 vg/g in mice. **E)** Protocol: wild-type neonatal Sprague Dawley rats (P4) received an AAV9 encoding TTL-dsRed at an intermediate dose (4.4E11 vg/g, N=3). After 6 weeks, single ventricular myocytes were isolated and functionally assessed. Following assessment, myocytes were binned into tertiles based on no/low, medium, or high level of TTL (indicated by dsRed fluorescence intensity). **F)** Myocyte stiffness (elastic modulus) measured via transverse nanoindentation. **G)** Sarcomere shortening representative traces and quantification of fractional shortening, contraction and relaxation velocity. **H)** Calcium transient representative traces and quantification of amplitude, rise time and decay time. Data are expressed as mean ± SEM with N/n representing the number of analysed mice and cardiomyocytes, respectively. Statistical significance was assessed via one-way ANOVA with post hoc Tukey test for multiple comparisons. Abbreviation: ns, non-significant.

We next evaluated isolated cardiomyocyte function upon TTL overexpression. We injected a single intermediate dose of AAV9-TTL-dsRed into P4 rats (4.4E11 vg/g), and isolated single cardiomyocytes 6 weeks later (Figure 1E). Rat myocytes were used for their robustness for functional characterization in our hands.^8, 9^ We took advantage of the heterogeneous expression of TTL-dsRed between cardiomyocytes from a given heart to compare myocyte properties to non-expressing controls from the same animal. Myocytes from transduced hearts could be binned into tertiles based on their TTL level (measured by dsRed intensity): 1) those expressing no/low levels of TTL (dsRed intensity within the range of uninjected myocytes, 52%), 2) those with moderate TTL expression (10-20x increase in dsRed level compared to control mean, 23%), and 3) those with high TTL overexpression (>20x increase in dsRed level, 27%).

Myocytes were assessed for changes in myocyte stiffness, intracellular calcium dynamics and unloaded shortening upon TTL overexpression. Following the functional assessment (blind to expression level), myocytes were binned into TTL expression tertiles. Myocytes expressing TTL showed a dose-dependent reduction in myocyte viscoelasticity as measured via transverse nanoindentation (Figure 1F). Consistent with reduced internal stiffness, myocytes showed a TTL dose-dependent increase in fractional shortening and contractile kinetics (Figure 1G), which appeared to be independent of any changes in intracellular calcium cycling (Figure 1H). This data reflects previous observations obtained in human failing and non-failing cardiomyocytes after acute (48 h) adenoviral-mediated TTL overexpression,^9^ and suggests that the rapid reduction in myocyte stiffness and improved contractility (independent of calcium transient alterations as previously shown^8^) persist with chronic TTL overexpression.

### Chronic TTL overexpression improves cardiac function in Mybpc3-targeted KI mice

Next, we evaluated whether chronic TTL application could rescue the cardiac disease phenotype of the HCM KI mice *in vivo*. KI mice developed LV hypertrophy combined with systolic and diastolic dysfunction.^10, 11^ AAV9-Empty or AAV9-TTL vectors were systemically delivered with a dose of 2.25E+11 vg/g in 3-week-old WT and KI mice (Figure S2B,C). Cardiac function was evaluated by serial echocardiography 1, 2, 4, 8, and 12 weeks after treatment (Table S2). After 12 weeks of treatment, ejection fraction (EF), stroke volume (SV), cardiac output (CO), fractional area change (FAC), and aortic ejection time (AET) were lower in KI-Empty than WT-Empty (Figure 2A-E), whereas left ventricular mass-to-body weight ratio (LVM/BW) was higher (Figure 2F). TTL therapy partially ameliorated EF, SV and CO, but not FAC, AET and LVM/BW in KI mice. Similar data were obtained for the ventricular weight-to-BW (VW/BW), and the heart weight-to-BW (HW/BW), VW/tibia length (TL) and HW/TL ratios (data not shown). Overall, most parameters did not change between KI-TTL and KI-Empty groups, but a trend towards better EF, SV and CO from 4 weeks on after treatment was visible (Table S2). When the difference in parameters were calculated between the last and first echocardiography, EF did not significantly differ between the groups (Figure 2G), whereas both SV and CO were significantly higher in KI-TTL than in KI-Empty mice (Figure 2H,I). Among the parameters of diastolic function, isovolumic relaxation time (IRVT) was prolonged in KI-empty mice but not rescued by TTL treatment (Table S2). When only female mice were considered, the difference between KI-TTL and KI-Empty mice was stronger, and some of the systolic and hypertrophic parameters were ameliorated, such as EF, SV, CO, VW/BW and VW/TL (Figure S3).

**Figure 2.**
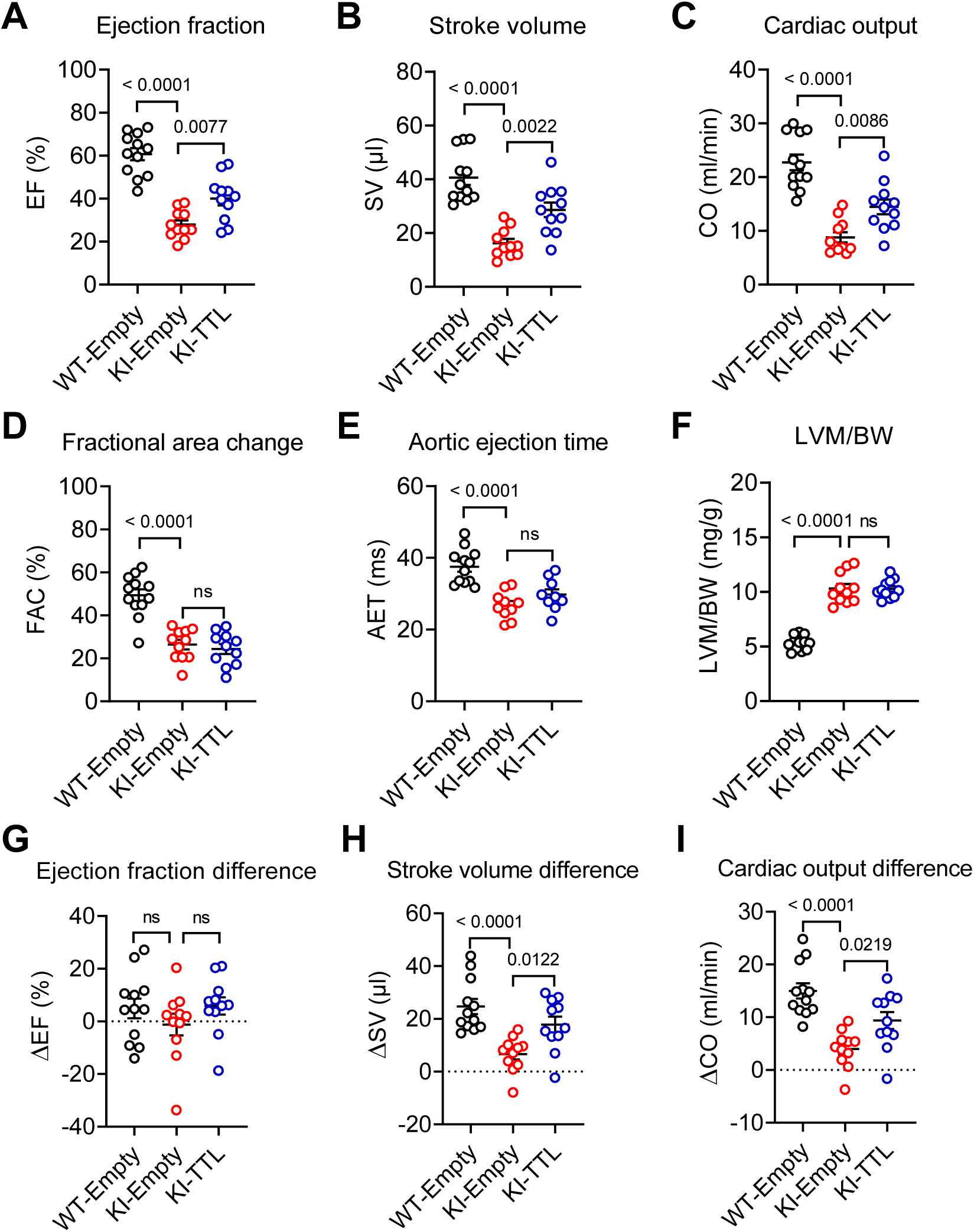
Evaluation of the cardiac phenotype by echocardiography after a 12-week AAV9-TTL/-Empty treatment. Three-week-old wild-type (WT) and *Mybpc3*-targeted knock-in (KI) mice were included in the study. Mice received either AAV9-Empty (no insert) or AAV9-TTL (HA-tagged human TTL). **A)** Ejection fraction (EF), **B)** Stroke volume (SV), **C)** Cardiac output (CO), **D)** Fractional area change (FAC), **E)** Aortic ejection time (AOT), **F)** Left ventricular mass to body weight ratio (LVM/BW). Difference in parameter values obtained between last and first echocardiography for **G)** Ejection fraction (ΔEF), **H)** Stroke volume (ΔSV) and **I)** Cardiac output (ΔCO). Data are expressed as mean ± SEM; Statistical significance was assessed via one-way ANOVA, followed by Dunnett’s post-test. Abbreviation: ns, non-significant.

At the end of the 12-week treatment, mice were subjected to hemodynamic measurements by LV catheterization (Figure 3). Global function was lower in KI-Empty than WT-Empty and TTL therapy restored SV, CO and stroke work back to WT levels in KI mice (Figure 3A-C, Table S3, P<0.05 with Student’s t-test). Systolic function was not affected, except for the lower EF and a trend to higher LV end-systolic volume (LVESV) and lower preload recruitable stroke work (PRSW) in KI-Empty mice (Figure 3D-H, Table S3). TTL did not impact on the systolic parameters (Figure 3D-H, Table S3). Diastolic function (dP/dt_min_ and Tau) was impaired in KI-Empty mice and not improved after TTL therapy (Figure 3I,L). On the other hand, whereas the LV end-diastolic volume (LVEDV) and minimal rate of volume change (dV/dt_min_) did not differ between KI- and WT-Empty, they were both higher after TTL therapy (Figure 3J,K), supporting the higher LV end-diastolic diameter (LVEDD) measured by echocardiography, which already increased after 2 weeks of TTL therapy (Table S2). PV loops of a representative mouse of each group at different stages of the occlusion showed higher LV pressure and lower SV in KI-Empty and a marked increase in LVEDV in KI-TTL mouse, resulting in a higher SV (Figure 3M,B,J). This was associated with a reduction in end-diastolic pressure-volume relation (EDPVR) in KI-TTL (p=0.0192 vs. KI-Empty, Student’s t-test; Figure 3N), suggesting reduced stiffness. Taken together, these data provide evidence that chronic TTL overexpression improved myocardial compliance and increased diastolic filling to restore CO in KI mice.

**Figure 3.**
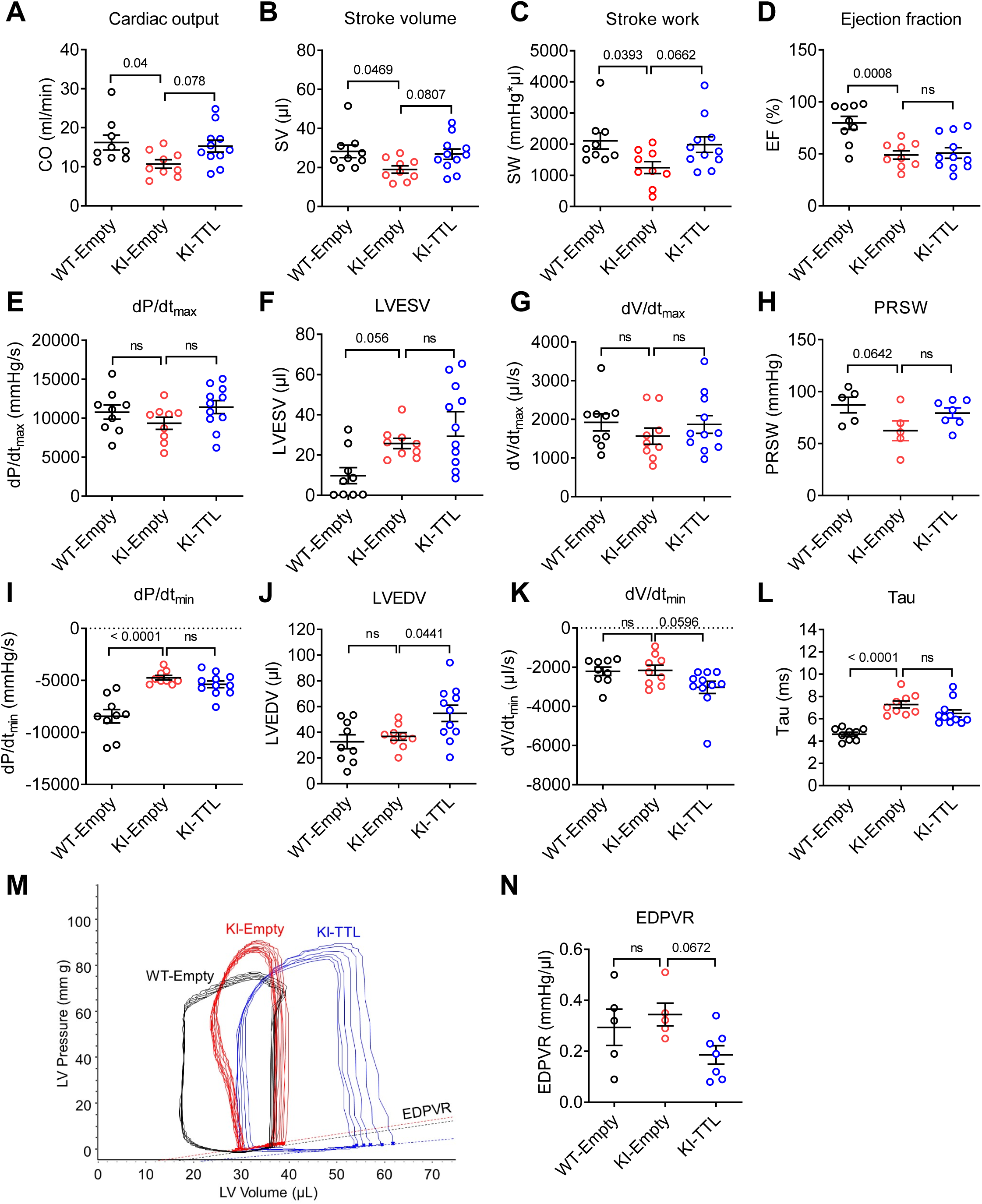
Evaluation of the cardiac phenotype by hemodynamics after a 12-week AAV9-TTL/-Empty treatment. Three-week-old wild-type (WT) and *Mybpc3*-targeted knock-in (KI) mice were included in the study (N=9-11). Mice received either AAV9-Empty (no insert) or AAV9-TTL (HA-tagged human TTL). **A)** Cardiac output (CO), **B)** Stroke volume (SV), **C)** Stroke work (SW), **D)** Ejection fraction (EF), **E)** Maximum rate of left ventricular pressure change (dP/dt_max_), **F)** Left ventricular end-systolic volume (LVESV), **G)** Maximal rate of left ventricular volume change (dV/dt_max_), **H)** Preload recruitable stroke work (PRSW), **I)** Minimal rate of left ventricular pressure change (dP/dt_min_), **J)** LV end-diastolic volume (LVEDV), **K)** Minimal rate of left ventricular volume change (dV/dt_min_), **L)** Time constant of active relaxation (Tau), calculated with the Weiss method, **M)** PV-loop traced after increasing occlusion in a representative mouse of each performed group, **N)** End-diastolic pressure-volume relation (EDPVR), calculated with a linear fit. Data are expressed as mean ± SEM. P values vs. KI-Empty were obtained with one-way ANOVA, followed by Dunnett’s multiple comparisons test. Abbreviation: N, number of mice; ns, non-significant.

### Chronic TTL overexpression reduces tubulin detyrosination in KI mice

TTL protein level did not differ between KI-Empty and WT-Empty and was 37-fold higher in cardiac cytosolic protein fraction of KI-TTL than in KI-Empty (Figure 4A,B). The levels of α-tubulin and dTyr-tub were higher in cytoskeletal-enriched protein fractions of KI-Empty than in WT-Empty (Figure 4C-E), suggesting increased stable microtubule network in KI mice. TTL did not impact α-tubulin, whereas it reduced dTyr-tub level in KI mice. The remaining dTyr could be due to the compensation by the up-regulation of other tubulin detyrosinases in KI-TTL (see below).

**Figure 4.**
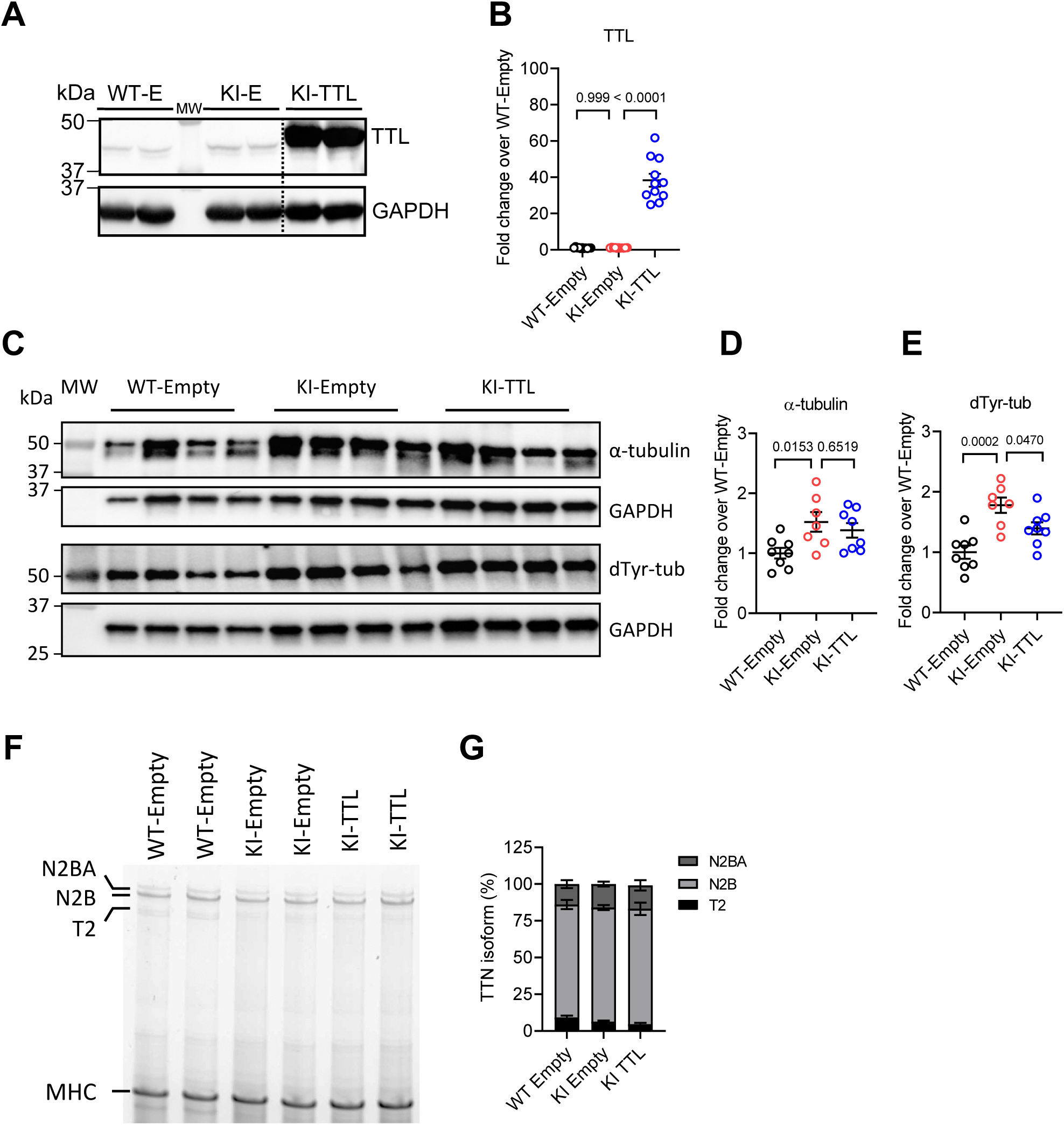
Molecular evaluation of tubulin detyrosination and titin isoforms in mice. **A)** Representative Western blot of cardiac cytosolic protein fractions stained for tubulin tyrosine ligase (TTL) and for GAPDH as a loading control in WT-Empty (WT-E), KI-Empty (KI-E) and KI-TTL mice; dashed line indicates where the blots were cut. **B)** TTL protein levels normalized to GAPDH (N=11). **C)** Representative Western blot of cardiac cytoskeletal-enriched protein fractions stained for α-tubulin, detyrosinated tubulin (dTyr-tub) and GAPDH as loading control. **D)** α-tubulin and **E)** dTyr-tub protein levels normalized to GAPDH (N=7-8). **F)** Representative Western blot of cardiac crude protein fractions stained for titin (isoforms N2BA, N2B and T2) and myosin heavy chain (MHC). **G)** Percentage of N2BA, N2B and T2 isoforms normalized to MHC (N=5, 8, 8 in WT-Empty, KI-Empty and KI-TTL, respectively). Data are expressed as mean ± SEM and relative to the mean of WT-Empty. P values were obtained with one-way ANOVA, followed by Tukey’s multiple comparisons test. Abbreviation: MW, molecular weight marker; N, number of mice.

### TTN isoform switch is not involved in decreased stiffness in KI-TTL

We then evaluated whether the reduction in EDPVR (Figure 3N) is the result of changes in titin isoform composition, leading to less N2B, stiff isoform, and more N2BA, compliant isoform, in KI-TTL. Long-run gel electrophoresis was performed as previously described^30^ in LV tissue extracted from the 3 mouse groups. No major difference was detected between the groups in the content of the 3 major isoforms of titin, N2BA, N2B and T2, and in the ratio of N2B/N2BA (Figure 4F,G). This suggests that titin isoform switch does not happen in KI mice and is not involved in the decreased stiffness in KI-TTL.

### Chronic TTL overexpression induces molecular changes in KI mice

To understand the molecular changes induced by long-term TTL overexpression, RNA-seq and mass spectrometry analyses were performed in female mouse LV tissue extracts (N=3 per group). Differentially expression analysis revealed 1013 dysregulated known mRNAs (263 higher, 750 lower, log2 ratio>0.58 or <-0.58, P<0.05) in KI-Empty than in WT-Empty, and only 104 dysregulated mRNAs (70 higher, 34 lower) in KI-TTL than in KI-Empty (Figure 5A,B). GO pathway analysis of RNA-seq data revealed enrichment (log2 ratio >0.58, P<0.05) in components of the sarcomere, actin cytoskeleton, T-tubule, L-type Ca^2+^ channel, microtubule, with enrichment of cellular and developmental process and transport in KI-Empty vs. WT-Empty (Figure S4A, dataset 1). Since the number of significantly higher and lower mRNAs in KI-TTL vs KI-Empty was low, GO analysis was done on all (log2 ratio >0.58 or <-0.58, P<0.05) and revealed dysregulated metabolic processes and components of mitochondrion, lysosome, nucleus, and late endosome membrane (Figure S4B, dataset 1). Volcano plot of MS analysis revealed much less dysregulated proteins in KI-Empty than in WT (37 higher, 14 lower, log2 ratio >0.58 or <-0.58, P<0.05; Figure 5C) and in KI-TTL than in KI-Empty (11 higher, 9 lower, log2 ratio >0.58 or <-0.58, P<0.05; Figure 5D). MS all pathway analysis in KI-Empty vs. WT-Empty (log2 ratio >0.58 or <-0.58, P<0.05) revealed alterations in components of the sarcomere, succinate-CoA ligase complex, intercalated disk, proteasome complex, and in muscle development and myofibril assembly (Figure S4C, dataset 2). MS all pathway analysis in KI-TTL vs. KI-Empty (log2 ratio >0.58 or <-0.58, P<0.05) revealed changes in components of mitochondria, connective tissue, Z-disc, ribosomal subunits, intercalated disc, and actin cytoskeleton (Figure S4D, dataset 2).

**Figure 5.**
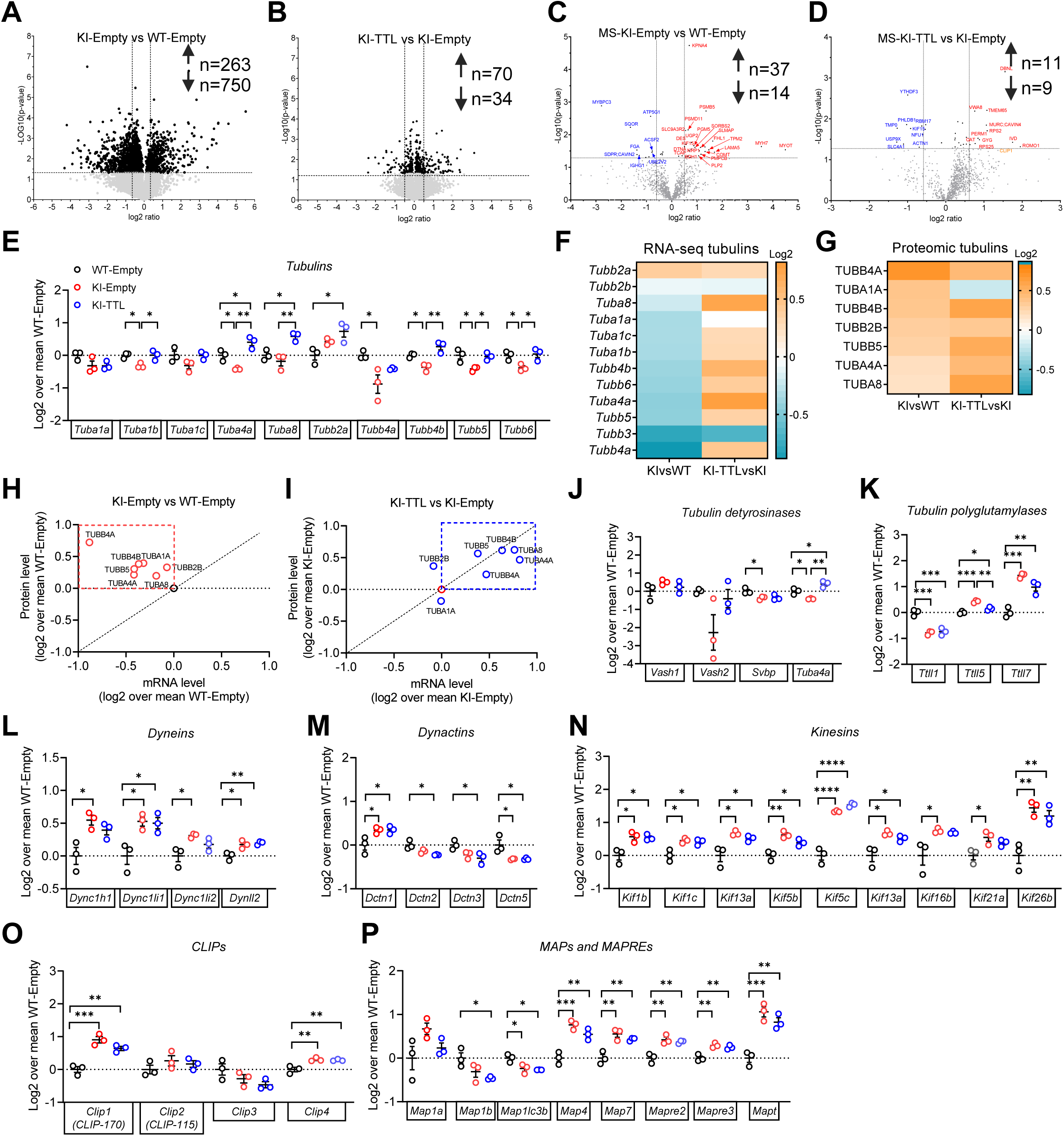
RNA-seq and proteome analysis in mouse hearts. RNA-seq and mass spectrometry analyses were performed in female mouse LV tissue extracts (N=3). Volcano plots show the -Log10 of P-value vs. the magnitude of change (log2 ratio) of **A)** mRNA levels in KI-Empty/WT-Empty, **B)** mRNA levels in KI-TTL/KI-Empty, **C)** protein levels in KI-Empty vs. WT-Empty and **D)** protein levels in KI-TTL vs. KI-Empty; light grey dots indicate P>0.05 and dark grey dots P<0.05. **E)** Tubulin mRNA levels over WT-Empty. **F)** Tubulin mRNA levels in KI-Empty/WT-Empty (left) and in KI-TTL/KI-Empty (right). **G)** Tubulin protein levels in KI-Empty/WT-Empty (left) and in KI-TTL/KI-Empty (right). Correlation of log2 ratio of tubulin mRNA and protein levels in **H)** KI-Empty/WT-Empty and **I)** KI-TTL/KI-Empty. **(J-P)** mRNA levels over WT-Empty of **J)** tubulin detyrosinases, **K)** tubulin polyglutamylases, **L)** dyneins, **M)** dynactins, **N)** kinesins, **O)** CAP-GLY domain containing linker proteins (CLIPs), **P)** microtubule-associated proteins (MAPs) and protein RP/EB family members (MAPREs). Data are expressed as log2 mean ± SEM over mean of WT-Empty or KI-Empty; *P<0.05, **P<0.01, ***P<0.001 and ****P<0.0001, one-way ANOVA, followed by Tukey’s post-test.

We then specifically analyzed the expression of tubulins and found lower levels of *Tuba1b*, *Tuba4a*, *Tubb5* and *Tubb6* mRNAs in KI-Empty, which were normalized with TTL treatment, and even higher levels of *Tuba4a*, *Tubb2a* and *Tuba8* in KI-TTL than in WT-Empty (Figure 5E). Heatmap revealed marked difference in log2 ratio of tubulin mRNAs between KI-Empty and KI-TTL (Figure 5F). These data were supported by the quantification of selected mRNAs using the nanoString nCounter® Elements technology and customized mouse-specific panels (Table S1). Several genes altered in heart failure were dysregulated in KI-Empty, such as *Myh7*, *Rcan1, Nppb*, *Fhl1*, *Myh6*, *Col1a1*, *Postn*, *Fhl2, Pln* and *Ppp1r1a* (Figure S4E). The levels of mRNAs encoding microtubule-associated proteins, such as *Mapt*, *Clip1*, *Mapre3, KiF5b*, *Map4*, *Dync1h1, Hdac6, Map1lc3a and Actr1a* were higher in KI-than in WT-Empty (Figure S4E). Similarly, the levels of mRNAs encoding proteins of the autophagy-lysosomal pathway such as *Bag3*, *Sqstm1*, *Jak1*, *Lamp1*, *Chmp2b,* and *Epg5* were higher in KI-Empty (Figure S4E), as shown previously in these mice.^37^ TTL respectively increased and decreased mRNA levels of several tubulin isoforms and α-tubulin deacetylase *Hdac6* in KI mice (Figure S4F).

Proteomic analysis of tubulins revealed less isoforms than in RNA-seq and an accumulation of all tubulin isoforms in KI- vs WT-Empty and to a higher extent in KI-TTL than in KI-Empty, except for TUBA1A (Figure 5G). Interestingly, tubulin mRNA and protein levels were inversely correlated in KI-Empty, i.e. lower mRNA levels were associated with higher protein levels than in WT-Empty (Figure 5H) and paralleled what was previously observed in heart failure samples.^29^ TTL reversed this correlation and induced the accumulation of most tubulin mRNAs and proteins (Figure 5I). This suggests that KI mice exhibit tubulin protein hyperstability, which could be related to longer lifetimes, increased autoinhibition of tubulin mRNAs, a new concept suggested recently.^29^ TTL drives an increase in both tubulin mRNA and protein levels, perhaps through autoactivation.

Whereas *Vash1* mRNA level did not differ between the groups, *Vash2* mRNA level was very variable in KI-Empty mice, and mRNA levels of both *Svbp* and *Tuba4a*, encoding a dTyr-tub isoform, were lower in KI-Empty (Figure 5J). *Tuba4a* level was 1.7-fold higher in KI mice treated with TTL (Figure 5E), suggesting that it could counteract the effect of TTL and therefore limit the reduction of dTyr-tub (Figure 4E). The mRNA levels of tubulin polyglutamylases were dysregulated in KI-Empty, and TTL almost completely normalized *Ttll5* that prefers α- over β- tubulins in KI mice (Figure 5K). Several mRNAs encoding other microtubule-associated components were altered in KI-Empty and were not affected by TTL therapy, such as the molecular motors dyneins, dynactins and kinesins (Figure 5L-N), CAP-Gly domain-containing linker proteins (CLIPs), microtubule-associated proteins (MAPs) and microtubule-associated protein RP/EB family members (MAPREs; Figure 5O,P).

Deeper proteome analysis revealed only 5 dysregulated proteins in KI mice that were normalized with TTL overexpression. This is the case for kinesin-12 (KIF15), the only kinesin isoform detected by proteomics, which was 1.75-fold higher in KI-Empty and fully normalized by TTL treatment (Figure S5A). Another one is the caveolae-associated protein 2 (CAVIN2/ SDPR; 1.5-fold lower), which interacts with CAVIN4 (MURC; also 1.5-fold lower) to facilitate the caveolae formation^38^ Similarly, some mitochondrial proteins were normalized by TTL, such as succinate-CoA ligase GDP/ADP-forming subunit alpha (SUCLG1; 1.3-fold lower in KI-Empty), NFU1 iron-sulfur cluster scaffold (NFU1; 1.5-fold higher), and electron transfer flavoprotein subunit beta (ETFB; 1.4-fold lower). RNAseq analysis showed that *SDPR*/*CAVIN2*, *SUCLG1 and ETFB* mRNA levels were also lower in KI-Empty than in WT-Empty (Figure S5B). However, none of them was normalized by TTL. KIF15 and NFU1 were not dysregulated at mRNA levels in KI-Empty and KI-TTL. The correlations between protein and mRNA levels in KI-Empty and KI-TTL vs WT-Empty shows that normalization by TTL was mainly on the protein levels (Figure S5C,D).

### TTL overexpression rescues hypertrophy in cardiomyocytes derived from a HCM hiPSC line

To translate our mouse data to humans, we used a HCM patient-derived heterozygous mutant *MYBPC3* (MYBPC3het; UKEi070-A) and its isogenic control (MYBPC3ic; UKEi070-A-1) hiPSC lines that were previously created.^12^ MYBPC3het carries the c.2308G>A genetic variant (p.Asp770Serfs98X), resulting in MYBPC3 protein haploinsufficiency in cardiomyocytes.^12^ We first evaluated the dose-response of AAV9-TTL in MYBPC3ic hiPSC-derived cardiomyocytes and found that a MOI of 100,000 gave rise to 19-fold higher TTL protein level than in non-transduced or AAV9-Empty-transduced cardiomyocytes (Figure 6A,B). We then evaluated the impact of a 10-day gene transfer of AAV9-TTL and AAV9-Empty (MOI 100,000) in both genotypes in the presence of H_2_O (Ctrl) or 100 nM endothelin-1 (ET1) for the last 3 days (Figure 6C-H), as previously described.^25, 39^ In basal condition, mean cell area was 1.6-fold higher in MYBPC3het than in MYBPC3ic, and TTL overexpression normalized cell hypertrophy in MYBPC3het (Figure 6C,D). ET1 induced cellular hypertrophy in MYBPC3ic and to a higher extent in MYBPC3het (3.4-fold higher than in MYBPC3ic), and AAV9-TTL normalized this effect in both groups (Figure 6F,G). These data revealed an anti-hypertrophic effect of TTL and activation of tubulin tyrosination in a human cellular model of HCM.

**Figure 6.**
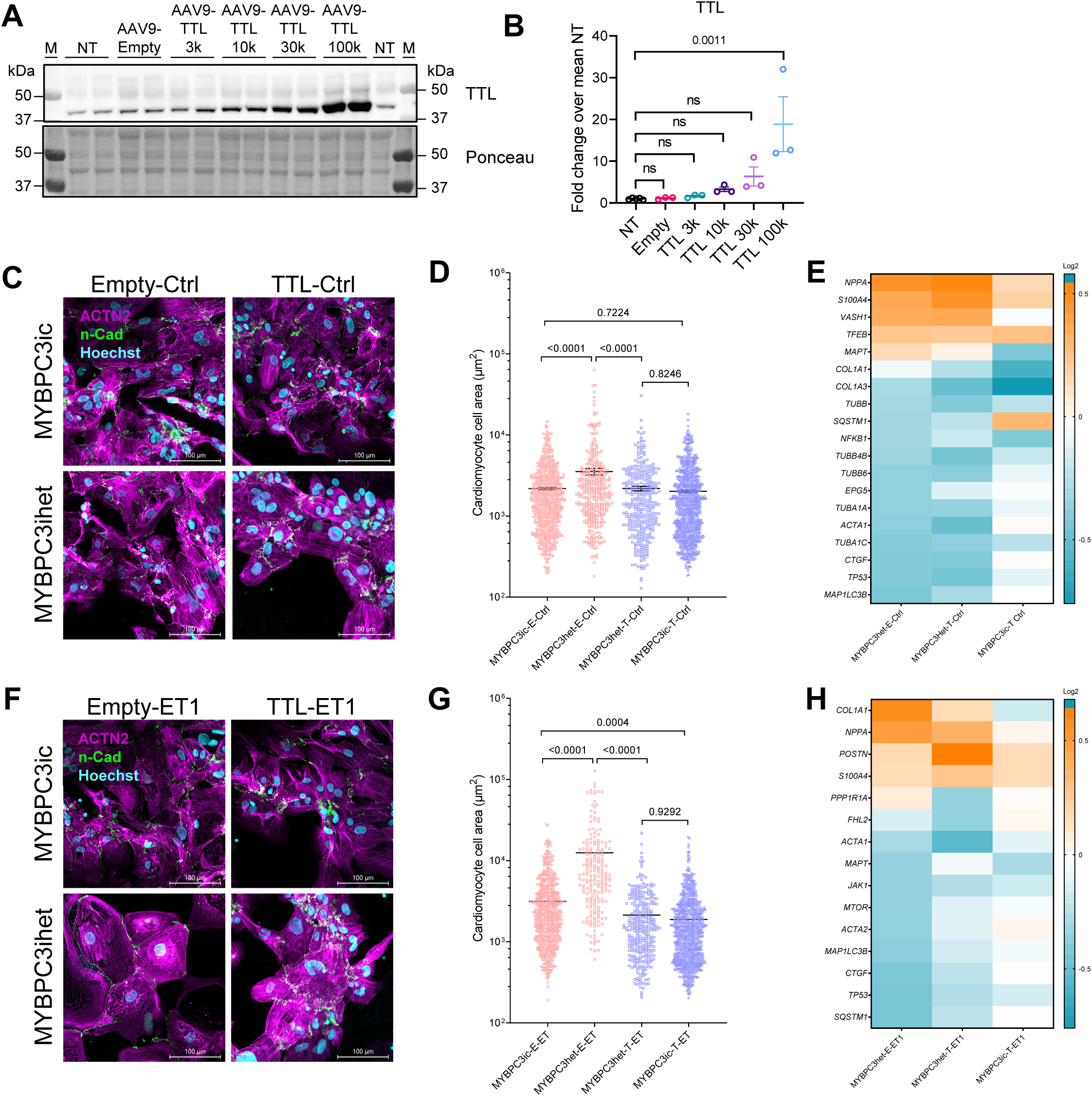
Evaluation of AAV9-mediated TTL transfer in MYBPC3ic and MYBPC3het hiPSC-derived cardiomyocytes. MYBPC3ic hiPSC line was differentiated into cardiomyocytes for 2 weeks, and then non-transduced (NT), transduced with AAV9-Empty (MOI 30 k), or transduced with AAV9-TTL (MOI 3 k to 100 k) for 7 days. **A)** Representative Western blot of MYBPC3ic crude protein fractions stained for TTL and respective Ponceau. **B)** Quantification of TTL protein levels, normalised to Ponceau and related to NT mean in MYBPC3ic hiPSC-CMs. MYBPC3ic and MYBPC3het hiPSC lines were differentiated into cardiomyocytes for 2 weeks, and then transduced with AAV9-Empty or AAV9-TTL (MOI 100 k) for 10 days in the presence of H_2_0 **(C-E)** or 100 nM endothelin-1 (ET1; **F-H**) for the last 3 days. Representative immunofluorescence analysis of MYBPC3ic and MYBPC3het hiPSC-cardiomyocytes transduced with AAV9-Empty or AAV9-TTL for 10 days in **(C)** basal condition or **(D)** after stimulation with ET1; hiPSC-CMs were stained for α-actinin 2 (ACTN2, purple), N-cadherin (N-Cad, green) and DAPI (blue). Scale bar = 100 µm. Quantification of cell area in hiPSC-CMs in **(D)** basal condition or **(G)** after stimulation with ET1 (results from 1 batch of differentiation). Selected hits of dysregulated RNA counts obtained with the nanostring analysis in **(E)** basal condition or **(H)** after stimulation with ET1 (results from 3-4 batches of differentiation). Data are expressed as mean ± SEM, with P values obtained with a one-way ANOVA, followed by a Tukey’s post-test. Abbreviations: AAV9, adeno-associated virus serotype 9; ns, non-significant.

All groups were subjected to a gene expression analysis using the nanoString nCounter® Elements technology and customized human-specific panels for heart failure, microtubules and autophagy. In basal condition, some markers were higher (*NPPA*, *S100A4*, *VASH1*, *TFEB*, *MAPT*), and markers or autophagy (*MAP1LC3B*, *SQSTM1*) and tubulin isoforms (*TUBA1C*, *TUBB* and *TUBB4B*, *TUBB6*) were lower in MYBPC3het than in MYBPC3ic, but TTL effect was not very clear (Figure 6E). In the presence of ET1, several markers were normalized in MYBPC3het with TTL, such as *COL1A1*, *NPPA*, *MAPT*, *MTOR*, *MAP1LC3B*, *SQSTM1* (Figure 6H).

### Chronic activation of tubulin tyrosination improves contractility in hiPSC-derived engineered heart tissues

To evaluate the impact of chronic tubulin tyrosination or detyrosination in a human context, hiPSC lines deficient in SVBP or TTL were created from a WT hiPSC line (mTag-RFP-T-TUBA1B) with CRISPR/Cas9 genetic tools (Figure S2A-D) and differentiated into cardiomyocytes. RT-qPCR confirmed the respective deficiency of SVBP and TTL mRNAs in hiPSC-cardiomyocytes (Figure S2E,F). SVBP-KO and TTL-KO hiPSC-derived cardiomyocytes respectively exhibited a markedly lower and higher dTyr-tub immunofluorescence intensity (Figure 7A) and protein levels (Figure 7B), confirming the targeting of either protein in hiPSC lines. The dTyr-tub protein level was not completely abolished in SVBP-KO, suggesting a compensatory role for TUBA4A and/or the recently discovered MATCAP1^40^ in this process.

**Figure 7.**
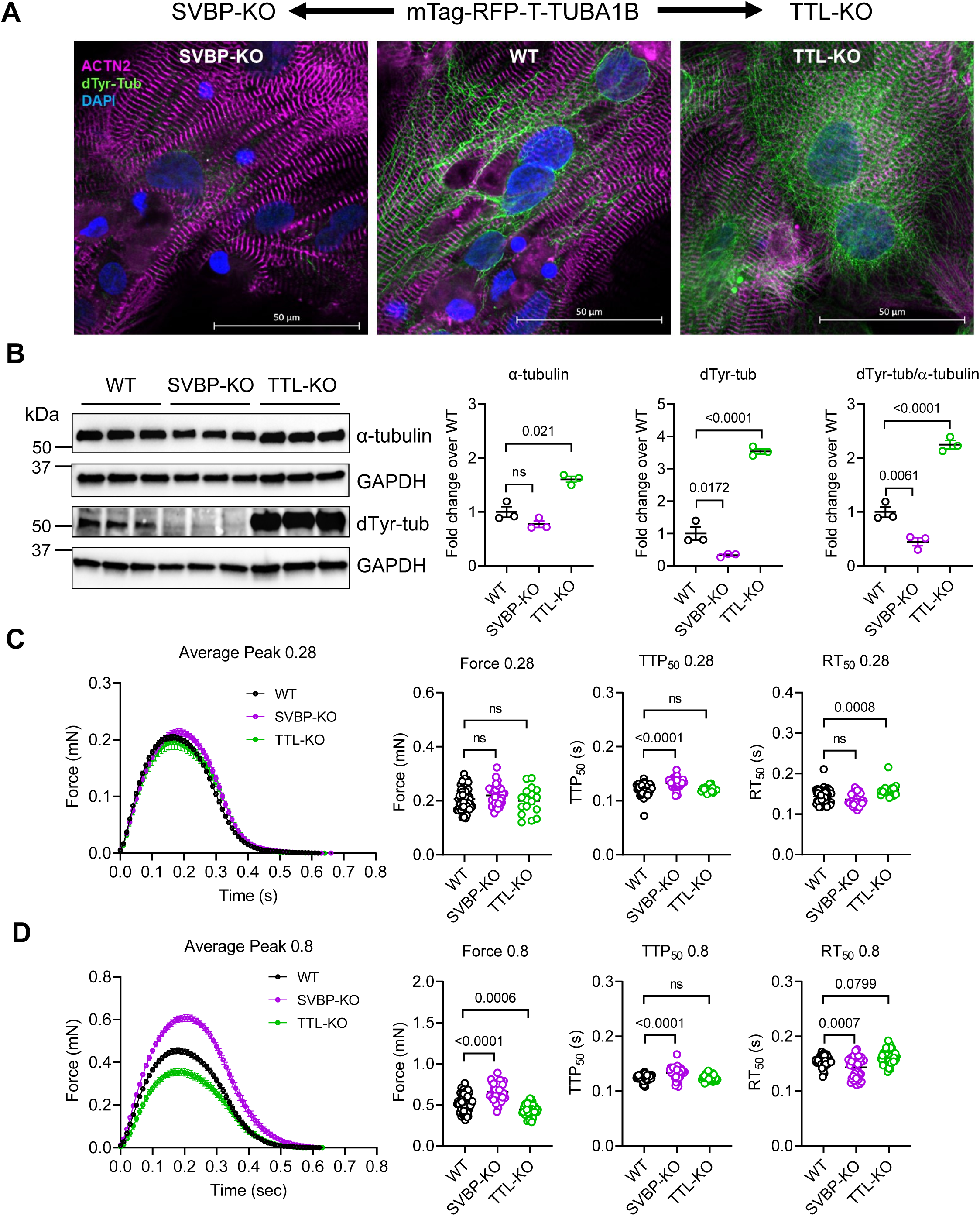
Evaluation of SVBP-KO and TTL-KO hiPSC-derived cardiomyocytes and engineered heart tissues. SVBP-KO and TTL-KO hiPSC lines were created from a control wild-type hiPSC line (WT, mTag-RFP-T-TUBA1B) with CRISPR/Cas9 genetic tools and differentiated into cardiomyocytes and engineered heart tissues (EHTs). **A)** Immunofluorescence analysis of 30-day-old SVBP-KO, WT and TTL-KO hiPSC-cardiomyocytes stained for α-actinin 2 (ACTN2, purple), detyrosinated α-tubulin (dTyr-tub, green) and DAPI (blue). Bar scale = 50 µm. **B)** Western blot was performed on 30-day-old hiPSC-cardiomyocytes (N=3) stained for α-tubulin, dTyr-tub and GAPDH, as a loading control. Quantification of α-tubulin, dTyr-tub and dTyr-tub/α-tubulin protein levels normalised to GAPDH and relative to WT mean. HiPSC-derived cardiomyocytes were cast in EHTs and cultivated for 60 days. Measurements of average peak, force amplitude, time to 50% of peak (TTP_50_) and time to 50% relaxation (RT_50_) of 60-day-old EHTs cast on **C)** standard posts (0.28 mN/mm; WT: N/d = 75/6, SVBP-KO: N/d = 33/5, TTL-KO: N/d = 17/2) or **D)** afterload enhanced posts (0.8 mN/mm; WT: N/d = 54/6, SVBP-KO: N/d = 33/5, TTL-KO: N/d = 38/5). Data are expressed as mean ± SEM, with P values vs. WT obtained with a one-way ANOVA, followed by a Dunnett’s post-test. Abbreviations: N/d, number of EHTs/differentiations; ns, non-significant.

Then, EHTs from the 3 genotypes were cast either on standard posts (0.28 mN/mm) or on stiffer posts that enhance afterload (0.8 mN/mm) and evaluated at the age of 60 days. On standard posts, force amplitude did not significantly differ between the groups, time to 50% peak (TTP_50_) was longer in SVBP-KO and did not differ to WT in TTL-KO (Figure 7C). Conversely, time from peak to 50% relaxation (RT_50_) was longer in TTL-KO and did not differ from the WT in SVBP-KO (Figure 7C). On stiffer posts, force amplitude and TTP_50_ were significantly higher, whereas RT_50_ was shorter in SVBP-KO (Figure 7D). In TTL-KO EHTs, force amplitude was lower, and kinetics were not significantly altered, but a trend towards longer RT_50_ was noticeable (Figure 7D). Therefore, stiffer posts were required to reveal an opposite phenotype in SVBP-KO and TTL-KO. These data provide evidence in a human cellular context that chronic tubulin tyrosination (=SVBP-KO) improved contractility in human EHTs, whereas chronic tubulin detyrosination (=TTL-KO) leads to contractility impairment.

RNA-seq and mass spectrometry analyses were then performed in EHTs from WT on standard posts and from WT, SVBP-KO and TTL-KO on stiff posts. GO analysis of RNA-seq data revealed that stiffer posts induced an enrichment (log2 ratio >0.58, p<0.05) in components of mitochondria and of metabolic process, suggesting maturation in WT EHTs (Figure S6A, dataset 3). SVBP-KO exhibited enrichment vs. TTL-KO in components of extracellular matrix, endoplasmic reticulum, Golgi, plasma membrane, and sarcomere, associated with an enrichment in developmental process, response to stimulus, actin filament-based process, regulation of proteolysis and muscle contraction (Figure S6B, dataset 3). Conversely, TTL-KO showed enrichment vs. SVBP-KO in components of mitochondria, ribosome, RNA-binding, polysome, nucleus, associated with enrichment in process of oxidative phosphorylation, mitochondrial electron transport, translation, mRNA degradation, and several other metabolic process (Figure S6C, dataset 3).

GO analysis of MS data showed enrichment in components of extracellular exosomes, vesicles and mitochondria in WT EHTs cast on stiffer posts (Figure S6D, dataset 4). SVBP-KO exhibited enrichment vs. TTL-KO in several cardiomyocyte components, including stress and contractile fibers, actin cytoskeleton, costamere, intracellular organelles, Z-disc and intercalated disc proteins, which were involved in cytoskeletal and actinin protein binding, actin filament-based process, muscle contraction, exocytosis and gluconeogenesis (Figure S6E, dataset 4). Conversely, GO analysis of TTL-KO showed enrichment vs. SVBP-KO in components of endoplasmic reticulum, Golgi, mitochondria, and secreted vesicles, which were involved in intracellular transport, metabolic process, negative regulation of proteolysis and response to stress (Figure S6F, dataset 4).

## Discussion

This study evaluated the impact of chronic activation of tubulin tyrosination in HCM mice, HCM hiPSC-cardiomyocytes and in hiPSC-EHTs. The main findings are: i) 6 weeks TTL overexpression dose-dependently reduced tubulin detyrosination and improved cardiomyocyte contractility without affecting calcium transients in WT rodent cardiomyocytes; ii) long-term AAV9-TTL overexpression improved diastolic filling, cardiac output and stroke volume and reduced stiffness without impact on TTN isoform composition in KI mice; iii) AAV9-TTL gene transfer for 10 days normalized cell hypertrophy in HCM hiPSC-cardiomyocytes; iv) TTL overexpression induced a marked transcription and translation of several tubulin isoforms and modulated mRNA or protein levels of components of mitochondria, sarcomere Z-disc, ribosomal subunits, intercalated disc, lysosome and cytoskeleton in KI mice; v) SVBP-deficient EHTs exhibited much lower dTyr-tub levels, higher force and faster relaxation than in TTL-deficient and WT EHTs under afterload; vi) RNA-seq and mass spectrometry analysis revealed distinct enrichment of cardiomyocyte components and pathways in SVBP-KO and TTL-KO EHTs.

KI mice exhibited lower EF, SV, CO and dP/dt_min_ and higher LVM/BW, HW/BW and HW/TL, supporting reduced systolic and diastolic function and LV hypertrophy, as shown previously.^10, 11, 17^ In addition, the expression of several genes involved in heart failure and autophagy-lysosomal pathway were dysregulated in KI mice, as reported previously.^37^ In this study, we also show that the mRNA levels of several microtubule-associated proteins, such as dyneins, dynactins, kinesins, CLIPs, MAPs and MAPREs are higher in KI than in WT mice. This suggests a broad up-regulation of a microtubule-transport program in KI mice. Proteomic analysis revealed higher abundance of sarcomeric, cytoskeletal and intercalated disc proteins, such as MYH7, DES, TCAP, FHL1, PGM5, XIRP1, and MYOT, and of components of the proteasome core complex, such as PSMA6, PSMB5 and PSMD11, supporting previous findings of increased proteasomal activity at this age.^41^. The level of dTyr-tub was higher in KI-than in WT-Empty as previously shown,^3^ A 6-week administration of AAV9-TTL dose-dependently reduced microtubule detyrosination in WT mice, supporting previous findings obtained acutely in human intact cardiomyocytes.^8, 9^ On the other hand, the elevated microtubule detyrosination was not fully normalized in *Mybpc3*-KI mice treated with TTL for 12 weeks. This could be explained by the accumulation of TUBA4A mRNA and protein, which is the only tubulin isoform translated as detyrosinated,^4, 7^ and of the detyrosinase MATCAP1,^40^ rather than a transcriptional upregulation of the VASH1/SVBP complex in KI-TTL mice.

A 6-week *in vivo* AAV9-TTL administration reduced stiffness and improved fractional shortening and velocities of contraction and relaxation in a dose-dependent manner in cardiomyocytes isolated from WT rats. In HCM KI mice, chronic TTL therapy ameliorated SV and CO, EF but not FAC, markedly increased diastolic filling and reduced stiffness. The latter was not associated with a shift of compliant-to-stiff TTN isoform in KI mice. On the other hand, TTL did not improve diastolic function (dP/dt_min_) and had no impact on LV hypertrophy, except for an amelioration of the VW/BW and VW/TL ratios in female KI-TTL mice. On the contrary, a 10-day TTL therapy normalized hypertrophy and partially corrected the hypertrophic transcriptome in human HCM cardiomyocytes. The positive impact of chronic TTL therapy on HCM mouse heart function was supported by the findings of markedly reduced level of dTyr-tub/α-tubulin, higher force of contraction and faster relaxation in SVBP-KO hiPSC-cardiomyocytes and -EHTs. Conversely, the TTL-KO cardiomyocytes and EHTs showed >2-fold higher levels of dTyr-tub/α-tubulin, lower force of contraction and prolonged relaxation. These data provide evidence in a human cellular context that chronic activation of tubulin tyrosination (=SVBP-KO) improved contractility in human EHTs, whereas chronic activation of tubulin detyrosination (=TTL-KO) induced contractility impairment, mimicking the situation in several types of heart failure^2, 3^ (for reviews,^4-7^).

Contractility findings in EHTs were supported by the RNA-seq and proteomic analyses, showing enrichment in components involved in muscle structure development, response to stimulus, cytoskeleton binding and function, and muscle contraction, whereas TTL-KO was rather enriched in components involved in oxidative phosphorylation, mitochondrial electron transport, translation, mRNA degradation, response to stress and other metabolic processes. Although RNA-seq and mass spectrometry data did not reveal the most likely candidates that could reduce stiffness in KI-TTL mice, GO pathway analysis revealed modulation of components of the cytoskeleton, mitochondria, metabolism, lysosome, sarcomere Z-disc and intercalated disc. TTL normalized protein, but not mRNA abundance of only 5 candidates (KIF15, SDPR, NFU1, SUCGL1 and ETFB). KIF15 (higher level in KI mice) has been shown to regulate tubulin polymerization in other contexts.^42^ SDPR/CAVIN2 (lower level in KI mice) interacts with CAVIN4/MURC to facilitate the SOCE and caveolae formation.^38, 43^ NFU1 (higher level in KI mice) is localized to mitochondria and plays a critical role in iron-sulfur cluster biogenesis. SUCGL1 (lower level in KI mice) is the ligase of succinate and CoA, leading to succinyl-CoA, which level was also found lower in patients with heart failure.^44^ ETFB (lower level in KI mice) interacts with connexin 43 in the heart and could participate in the mitochondrial redox state.^45^ We also found that CLIP1 protein level is further increased with TTL in KI mice, supporting previous findings that CLIP1 binds to the +end of Tyr-tub and improves dynein-dynactin-mediated retrograde cargo transport.^46, 47^ Another interesting finding in our study is that most of tubulin isoforms exhibited lower mRNA and higher protein level in KI mice, such as observed in human heart failure samples.^29^ In both models, TUBA1A, TUBB4B, TUBB2B and TUBB5 are more abundant, while TUBA1C, TUBB3 and TUBB6 proteins were not detected in our proteomic analysis. These data suggest that tubulin autoregulation, characterized by mRNA repression by abundant protein levels is effective in KI mice with hypertrophy and dysfunction, in agreement with previous work in human failing cardiomyocytes.^29^ Interestingly, however, TTL did not reverse this phenotype, but stimulated transcription of most tubulin isoforms, leading to accumulation of both tubulin mRNAs and proteins. This suggests that TTL-bound tubulin protein is not able to trigger autoinhibition as it should, leading to tubulin mRNA autoactivation.

Our study supports previous findings obtained after acute TTL treatment in isolated intact cardiomyocytes,^8, 9^ and provides evidence that a chronic AAV9-TTL application ameliorates diastolic filling and global cardiac function in HCM mice and normalized cellular hypertrophy in HCM hiPSC-cardiomyocytes. AAV9-based therapy has been tested in mouse or human cellular models of HCM (*MYBPC3*) and arrhythmogenic cardiomyopathy (*PKP2*). Long-term AAV9-*MYBPC3* therapy prevented the development of the cardiomyopathic phenotype in *Mybpc3*-deficient mice^17^ and normalized cellular hypertrophy in HCM hiPSC-cardiomyocytes^48^.^48^ Similarly, *PKP2* gene therapy reduced ventricular arrythmias, reverse remodeling of the right ventricle, improve heart function and extend survival in a *Pkp2*-deficient mouse model of arrhythmogenic cardiomyopathy.^49^ We are aware that AAV9 also targets the liver and could induce some liver toxicity,^50^ and the combination of AAV9 and a CMV promoter resulted in a lower detection of GFP in the mouse liver,^16^ On the other hand, the combination of AAV9 and the human cardiomyocyte-specific promoter *TNNT2* limits the expression of the transgene to the cardiomyocytes and not in the liver in mice.^17^ There are currently no results of clinical trials using AAV9-mediated gene therapy for cardiomyopathies, although a Phase 1b with AAV9-*MYBPC3* for HCM patients (NTC05836259) and a Phase 2 with AAV9-*LAMP2B* for patients with Danon disease (NTC06092034) have started.

In conclusion, this study provides the first proof-of-concept that chronic activation of tubulin tyrosination in HCM mice and in human EHTs improves heart function and holds promise for targeting the non-sarcomeric cytoskeleton in heart disease.

## Acknowledgments

The authors gratefully acknowledge Moritz Meyer-Jens and Eslem Nur Yueruemez (Pharmacology, UKE, Hamburg) for contributing to the maintenance of hiPSCs, EHTs and to experiments in hiPSC-derived cardiomyocytes, and the FACS core facility (UKE, Hamburg). We thank Zubayda Sultan and Jennifer Sarikaya (Amsterdam UMC) for the help in titin isoform analysis. We also would like to thank Thomas Eschenhagen and Marc Hirt (Pharmacology, UKE, Hamburg) for providing us with the standard and stiff silicone EHT posts. Finally, we thank Marie-Jo Moutin (GIN, Grenoble) for providing us with antibodies directed against tubulin modifications.

## Data Availability Statement

Datasets, analysis and study materials will be made available on request to other researchers for the purpose of reproducing the results or replicating the procedures. All data of OMIC experiments have been or will be made publicly available. The RNA-seq data have been deposited to RNA-seq datasets to NCBI’s Gene Expression Omnibus (GEO) of the European Nucleotide Archive (ENA) at EMBL-EBI, accession number is pending. The mass spectrometry proteomics data have been deposited to the ProteomeXchange Consortium via the PRIDE^51^ partner repository with the dataset identifier PXD042418.

