## Supplemental Figures and Tables for "Chronic activation of tubulin tyrosination in HCM mice and in human iPSC-engineered heart tissues improves heart function"

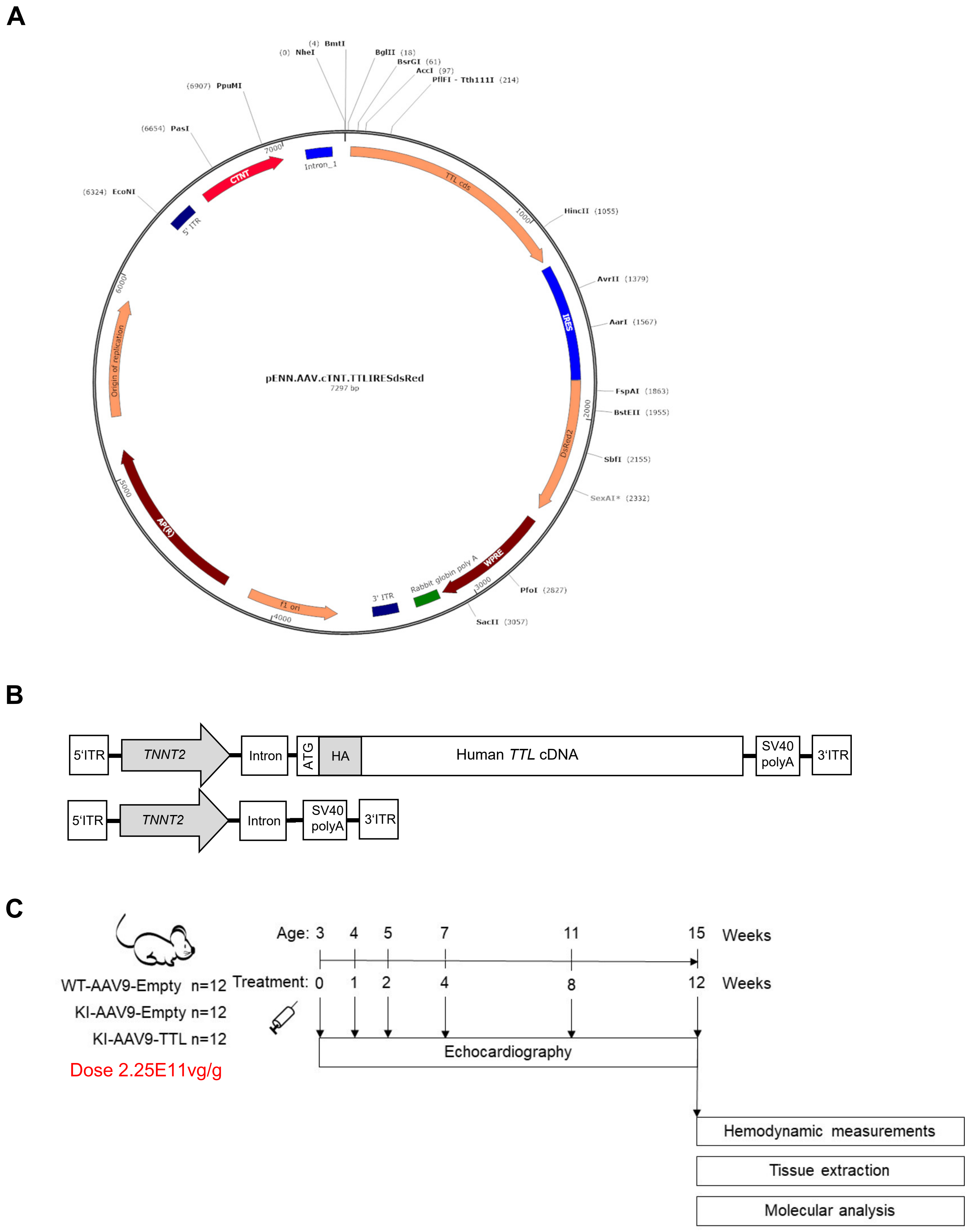

**Figure S1. Schematic representation of the plasmids used and protocol for TTL gene transfer in *Mybpc3*-targeted knock-in (KI) and wild-type (WT) mice. A)** Plasmid map of TTL-IRES-dsRed under the control of the human cardiac troponin T (cTnT; *TNNT2*) promoter (PENN.AAV.cTNT.TTL.IRES.dsRed). **B)** Schematic linear representation of the vector expressing the HA-tagged human WT *TTL* cDNA (AAV9-TTL; upper part) or nothing (AAV9-Empty; lower part) under the control of *TNNT2* promoter. **C)** Experimental protocol of AAV9-mediated gene transfer in mice, including the number of *Mybpc3*-targeted KI and WT mice used and the timeline of experiments.

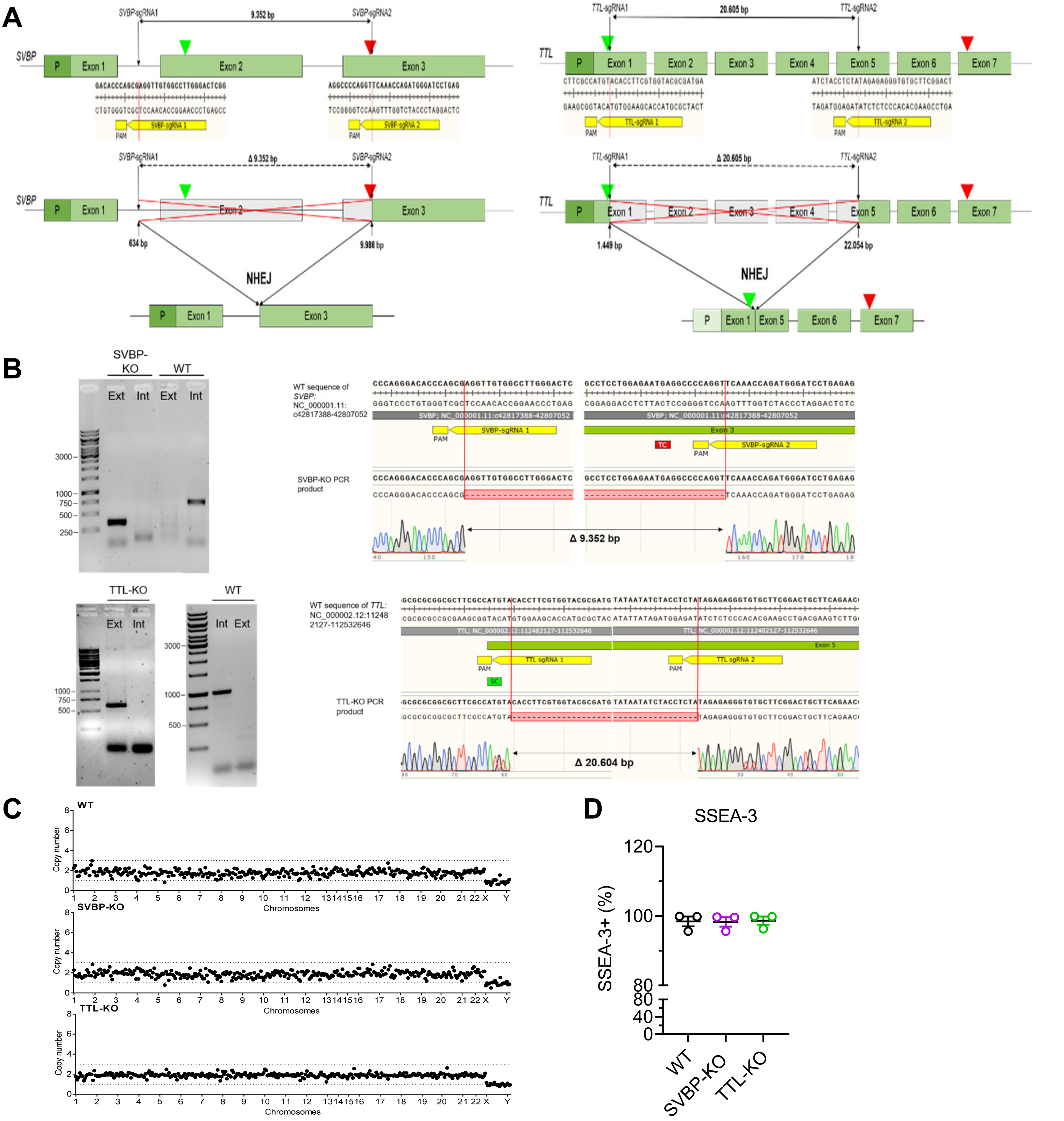

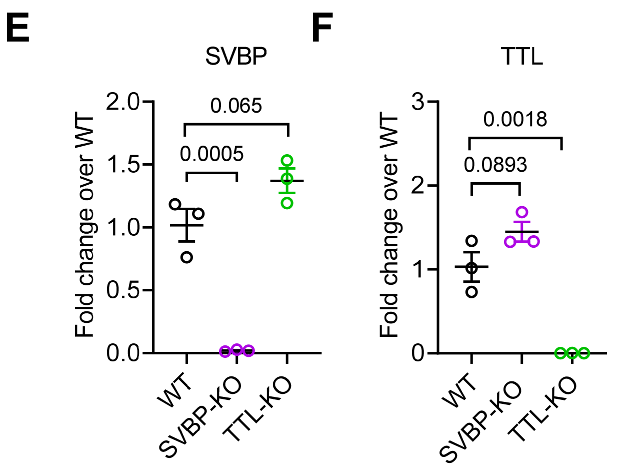

**Figure S2. Generation and characterization of SVBP-KO and TTL-KO hiPSC lines. A)** Schematic of knock-out (KO) approach for SVBP (left) and TTL (right) panel. **B)** Sequencing of KO DNA clones. PCR fragments for amplicons internal (Int) and external (Ext) of the large deletion are shown for SVBP-KO and WT DNA (upper) and for TTL-KO and WT DNA (lower). Alignment of the sequencing result for external fragments of the KO clones are shown on the right, confirming large, homozygous deletions (9,352 bp for SVBP-KO and 20,604 bp for TTL-KO. **C)** Nanostring karyotype panel analysis for WT and KO cell lines. **D)** Measurement of pluripotency by flow-cytometry SSEA-3 staining of hiPSC cultures. RT-qPCR results for **E)** *SVBP* and **F)** *TTL* mRNA levels in 30-day-old WT and KO hiPSC-derived cardiomyocytes. Data are expressed as mean ± SEM as fold change over mean WT, with P values obtained with one-way ANOVA, followed by Dunnett's multiple comparisons test.

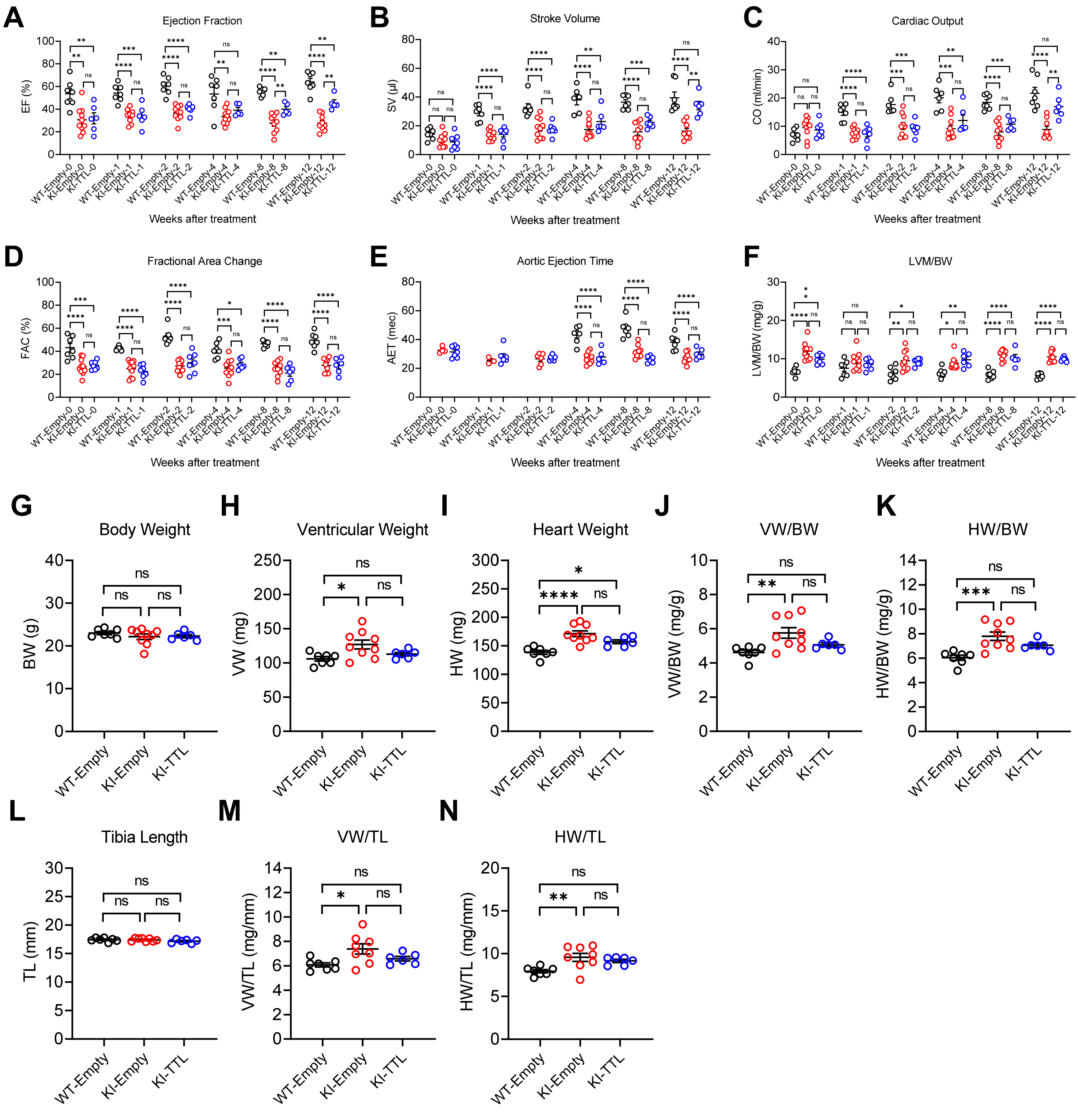

**Figure S3. Evaluation of the cardiac phenotype by echocardiography overtime and tissue weights in female mice.** Three-week-old wild-type (WT) and *Mybpc3*-targeted knock-in (KI) mice were included in the study. Mice received either AAV9-Empty (no insert) or AAV9-TTL (HA-tagged human tubulin tyrosine ligase). **(A-F)** Selected echocardiographic parameters overtime for **A)** Ejection fraction (EF), **B)** Stroke volume (SV), **C)** Cardiac output (CO), **D)** Fractional area change (FAC), **E)** Aortic ejection time (AET) and **F)** Left ventricular mass to body weight ratio (LVM/BW). **(G-N)** Mouse cardiac tissue weights and relation to body weight (BW) or tibia length (TL): **G)** BW, **H)** Ventricular weight (VW), **I)** Heart weight (HW), **J)** VW/BW, **K)** HW/BW, **L)** TL, **M)** VW/TL and **N)** HW/TL. Data are expressed as mean ± SEM, with *P<0.05, **P<0.01, ***P<0.001 and ****P<0.0001, one-way ANOVA, followed by Tukey's multiple comparisons test. Abbreviation: ns, non-significant..

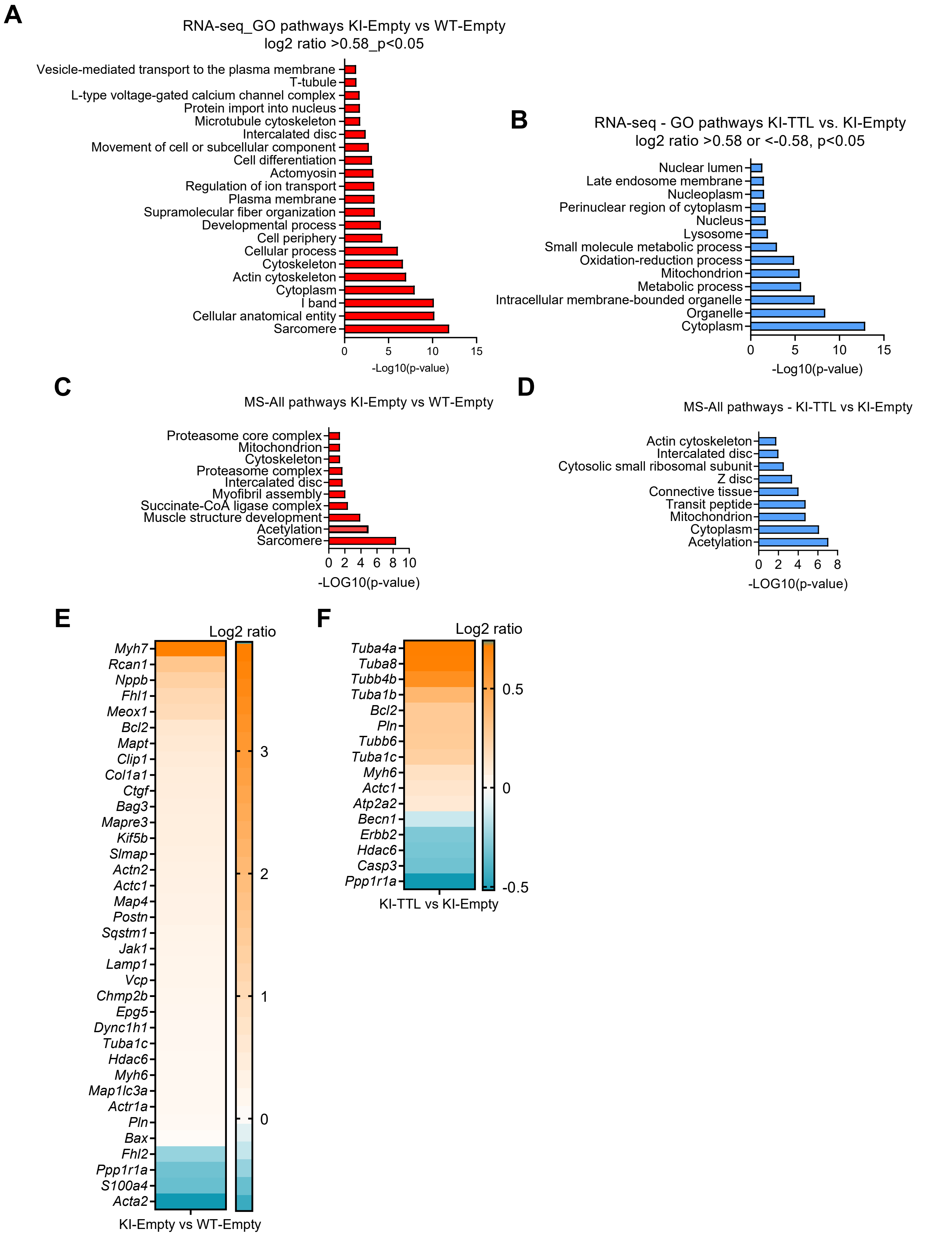

**Figure S4. GO pathway enrichment analysis in RNA-seq and mass spectrometry and nanostring analysis in mice.** Selected hits of GO pathways enrichment based on RNA-seq in **A)** KI-Empty vs. WT-Empty (log2 ratio>0.58, P<0.05, N=3) **B)** KI-TTL vs. KI-Empty (log2 ratio>0.58 or <-0.58, P<0.05, N=3). Selected hits of ALL string pathways based on MS analysis in **C)** KI-Empty vs. WT-Empty (log2 ratio >0.58 or <-0.58, p<0.05) and in **D)** in KI-TTL vs. KI-Empty (log2 ratio >0.58 or <-0.58, p<0.05). Quantification of levels of mRNAs encoding protein dysregulated in heart failure or involved in microtubules and autophagy using the nanostring nCounter Elements technology and customized mouse-specific panels, with **E)** log2 ratio in KI-Empty over mean of WT-Empty and **F)** log2 ratio of KI-TTL over KI-Empty (p<0.05; N=7 (WT), 9 (KI) and 6 (KI-TTL)). Only females were considered.

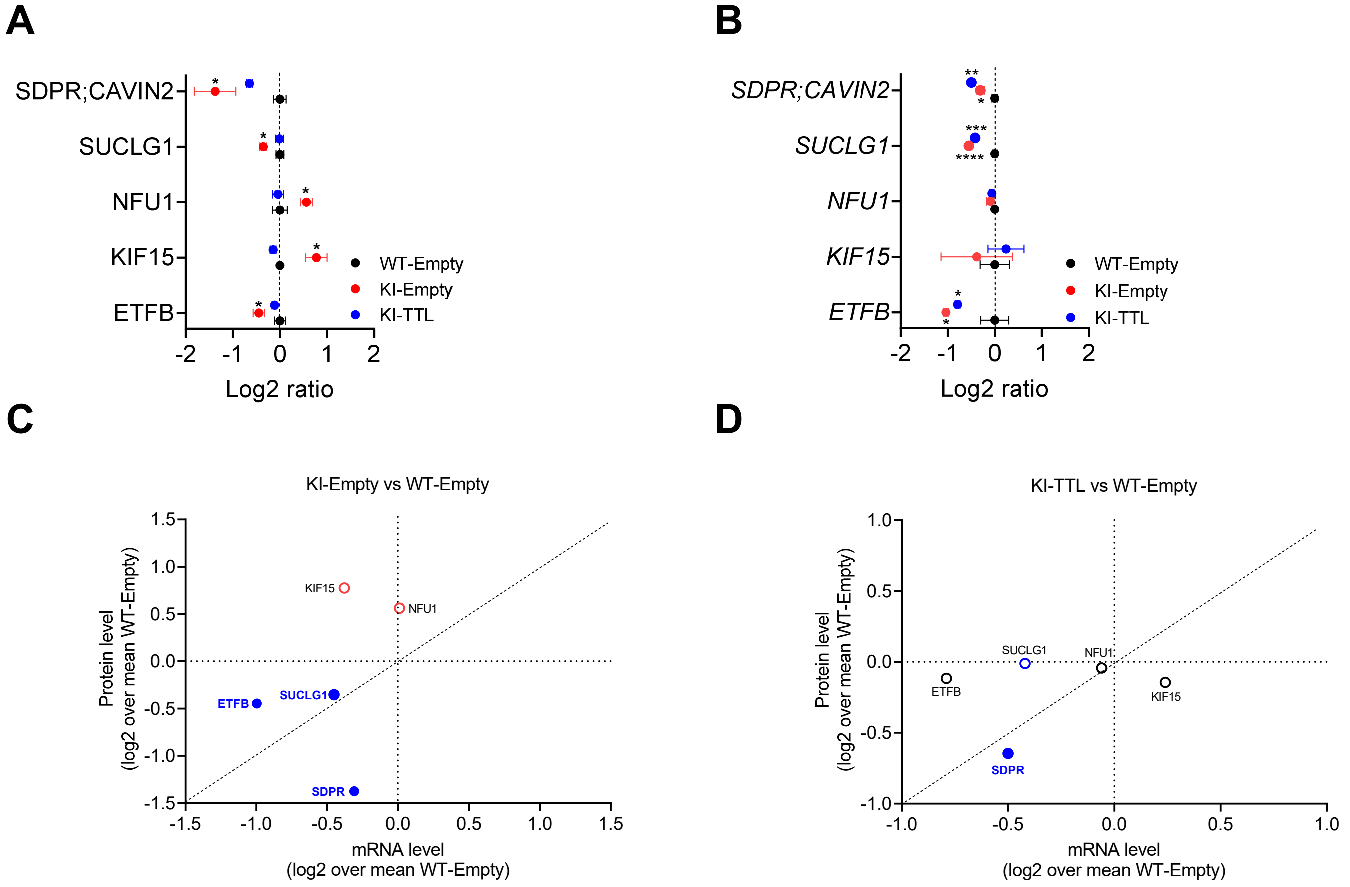

**Figure S5. Selected proteome and transcriptome analysis in mice and EHTs. A)** Significantly dysregulated proteins in KI-Empty vs mean WT-Empty and normalized by TTL. **B)** Corresponding mRNA levels over mean WT-Empty. **C)** Correlation of dysregulated proteins with their mRNA levels in KI-Empty vs WT-Empty; full symbols are significantly lower at both mRNA and protein levels in KI-Empty. **D)** Correlation of proteins and mRNA levels in KI-TTL vs WT-Empty; full symbol are significantly lower on both mRNA and protein levels in KI-TTL. Data are expressed as log2 mean ± SEM over mean of WT-Empty; *P<0.05, **P<0.01, ***P<0.001 and ****P<0.0001 vs. WT-Empty, one-way ANOVA, followed by Dunnett’s post-test.

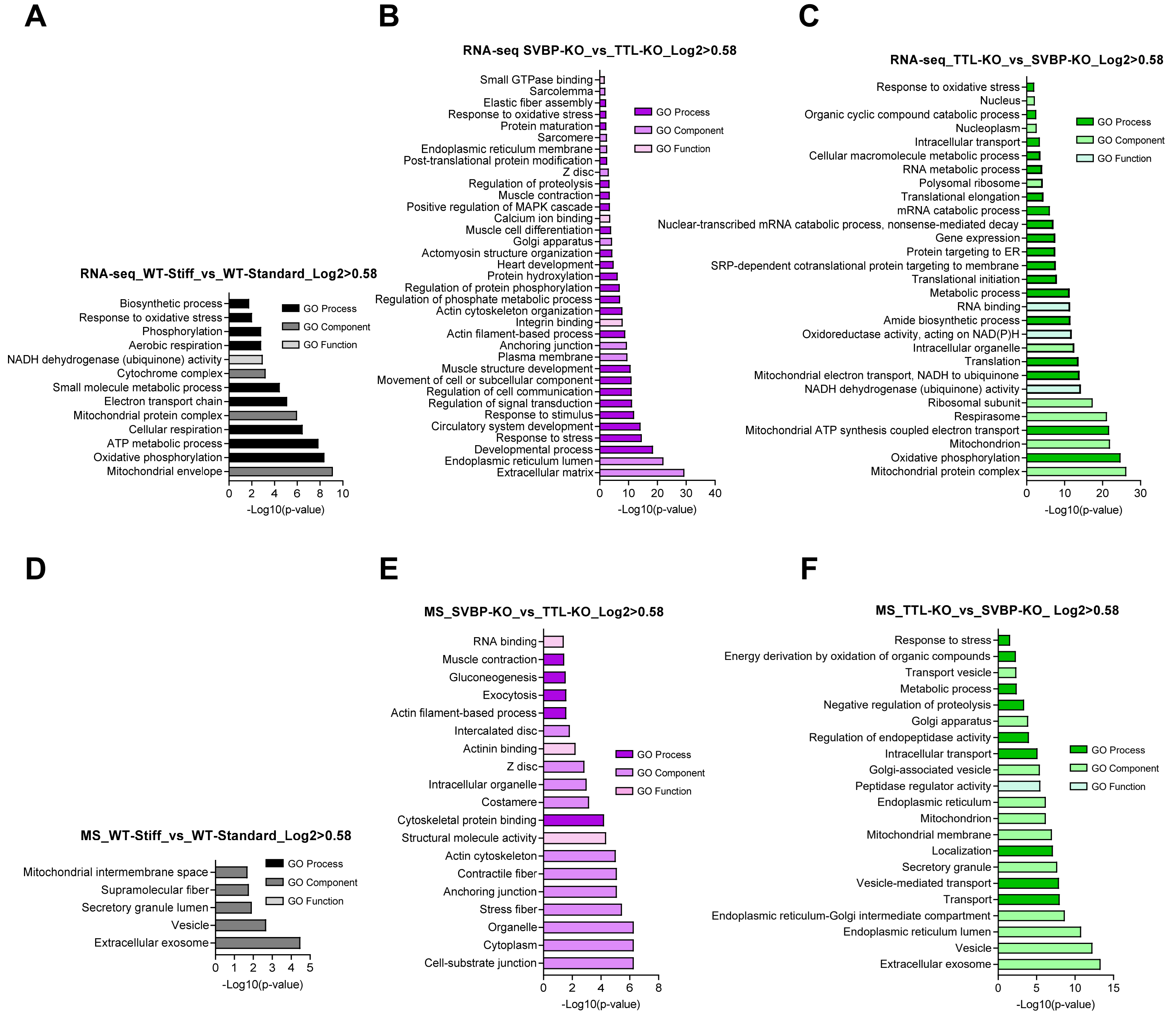

**Figure S6. GO pathway enrichment analysis in RNA-seq and mass spectrometry in hiPSC-derived engineered heart tissues (EHTs).** Selected hits of significantly enriched GO pathways based on RNA-seq data in **A)** WT-EHTs cast on stiff vs. standard posts (log2 ratio>0.58, P<0.05, N=3), **B)** SVBP-KO vs. TTL-KO cast on stiff posts (log2 ratio>0.58, P<0.05, N=2 vs. 3) and **C)** TTL-KO vs. SVBP-KO cast on stiff posts (log2 ratio>0.58, P<0.05, N=3 vs. 2). Selected hits of significantly enriched GO pathways based on mass spectrometry data in **D)** WT-EHTs cast on stiff vs. standard posts (log2 ratio>0.58, P<0.05, N=3), **E)** SVBP-KO vs. TTL-KO cast on stiff posts (log2 ratio>0.58, P<0.05, N=3) and **F)** TTL-KO vs. SVBP-KO cast on stiff posts (log2 ratio>0.58, P<0.05, N=3). Colours in bars correspond to the different GO pathways.

**Table S1. Customized NanoString's nCounter® Elements TagSet panels for mouse or human RNA analysis**

**Heart failure panel**

| **Acronym** | **Full name** |
|---|---|
| *Abcf1* | ATP binding cassette subfamily F member 1 |
| *Actb* | Actin, beta |
| *Acta1* | Actin, alpha, skeletal muscle |
| *Actc1* | Actin, alpha, cardiac muscle 1 |
| *Actn2* | Actinin alpha 2 |
| *Atp2a2* | ATPase sarcoplasmic/endoplasmic reticulum Ca2+ transporting 2 |
| *Bax* | BCL2 associated X, apoptosis regulator |
| *Bcl2* | BCL2, apoptosis regulator |
| *Casp3* | Caspase 3 |
| *Casq2* | Calsequestrin-2 |
| *Cdh5* | Cadherin 5, type 2, VE-cadherin |
| *Cltc* | Clathrin heavy chain |
| *Col1a1* | Collagen type I alpha 1 |
| *Col3a1* | Collagen type III alpha 1 |
| *Ctgf* | Connective tissue growth factor |
| *Fhl1* | Four-and-a-half-LIM-domains 1 |
| *Fhl2* | Four-and-a-half-LIM-domains 2 |
| *Gapdh* | Glyceraldehyde-3-phosphate dehydrogenase |
| *Meox1* | Mesenchyme homeobox 1 |
| *Myh6* | Myosin heavy chain 6 |
| *Myh7* | Myosin heavy chain 7 |
| *Nfkb1* | nuclear factor kappa B subunit 1 |
| *Nppa* | Natriuretic peptide A |
| *Nppb* | Natriuretic peptide B |
| *Pgk1* | Phosphoglycerate kinase 1 |
| *Pln* | Phospholamban |
| *Postn* | Periostin |
| *Ppp1r1a* | Protein phosphatase 1, regulatory (inhibitor) subunit 1A (PP-I-1) |
| *Rcan1* | Regulator of calcineurin 1 |
| *Ryr2* | Ryanodine receptor 2 |
| *S100a4* | S100 calcium binding protein A4 (=FSP1) |
| *Srf* | Serum response factor |
| *Vwf* | von Willebrand factor |

**Autophagy panel**

| **Acronym** | **Full name** |
|---|---|
| *Bag3* | BCL2 associated athanogene 3 |
| *Becn1* | Beclin-1 |
| *Chmp2b* | Charged multivesicular body protein 2B |
| *Epg5* | Ectopic P-granules autophagy protein 5 homolog |
| *Erbb2* | Erb-b2 receptor tyrosine kinase 2 |
| *Foxo1* | Forkhead box O1 |
| *Fyco1* | FYVE and coiled-coil domain containing 1 |
| *Hdac6* | Histone deacetylase 6 |

| | |
|---|---|
| *Jak1* | Janus kinase 1 |
| *Lamp1* | Lysosomal-associated membrane protein-1 |
| *Lamp2* | Lysosomal-associated membrane protein-2 |
| *Map1lc3a* | Microtubule associated protein 1 light chain 3 alpha |
| *Map1lc3b* | Microtubule associated protein 1 light chain 3 beta |
| *Mtor* | Mechanistic target of rapamycin |
| *Nbr1* | Neighbor of BRCA1 gene 1 |
| *Nfkb1* | Neuregulin 1 |
| *Rab7* | RAB7A; RAS oncogene family |
| *Sh3tc2* | SH3 domain and tetratricopeptide repeats 2 |
| *Sqstm1* | sequestosome 1 ; p62 |
| *Stat1* | Signal transducer and activator of transcription 1 |
| *Stat3* | Signal transducer and activator of transcription 3 |
| *Tfeb* | Transcription factor EB |
| *Trp53* | Tumor protein p53 |
| *Vcp* | Valosin containing protein (p97) |

**Microtubule panel**

| **Acronym** | **Full name** |
|---|---|
| *Actr1a* | ARP1 actin related protein 1 homolog A |
| *Clip1* | CAP-Gly domain containing linker protein 1 (CLIP-170) |
| *Clip2* | CAP-GLY domain containing linker protein 2 (CLIP-115) |
| *Dync1h1* | Dynein cytoplasmic 1 heavy chain 1 |
| *Kif2a* | Kinesin family member 2A (KHC, kinesin family 13) |
| *Kif5b* | Kinesin family member 5B (KHC, kinesin family 1) |
| *Map4* | Microtubule associated protein 4 |
| *Mapre1* | Microtubule associated protein RP/EB family member 1 (EB1) |
| *Mapre3* | Microtubule associated protein RP/EB family member 3 (EB3) |
| *Mapt* | Microtubule associated protein tau |
| *Slmap* | Sarcolemma associated protein |
| *Svbp* | Small vasohibin-binding protein |
| *Ttl* | Tubulin tyrosine ligase |
| *Tuba1a* | Tubulin alpha 1a |
| *Tuba1b* | Tubulin alpha 1b |
| *Tuba1c* | Tubulin alpha 1c |
| *Tuba4a* | Tubulin alpha 4a |
| *Tuba8* | Tubulin alpha 8 |
| *Tubb4b* | Tubulin beta 4B class Ivb (TUBB2C) |
| *Tubb5* | Tubulin, beta 5 class I |
| *Tubb6* | Tubulin beta 6 class V |
| *Vash1* | Vasohibin 1 |

**Table S2. Echocardiographic parameters overtime in *Mybpc3*-targeted knock-in (KI) and wild-type (WT) mice treated with AAV9-TTL and/or AAV9-Emtpy.**

| **Baseline / 3-week-old** | | | |
|---|---|---|---|
| Parameters | WT-Empty | KI-Empty | KI-TTL |
| N number | 12 | 12 | 12 |
| F/M | 7/5 | 12/2 | 7/5 |
| BW (g) | 8.9±0.2 | 7.6±0.3** | 8.3±0.3 |
| IVSd (mm) | 0.39±0.03 | 0.46±0.03 | 0.45±0.03 |
| LVPWd (mm) | 0.38±0.02 | 0.46±0.03 | 0.50±0.02** |
| LVEDD (mm) | 3.21±0.06 | 3.50±0.10* | 3.61±0.06*** |
| LVESD (mm) | 2.41±0.07 | 3.00±0.07**** | 3.13±0.06**** |
| FAC (%) | 44.0±2.7 | 25.1±2.1**** | 25.1±1.6**** |
| AET (msec) | nd | 32.6±1.1 | 31.4±1.5 |
| MV Decel Time (msec) | nd | 35.5±5.3 | 35.5±4.8 |
| MV E/A | nd | 0.95±0.06 | 1.21±0.08 |
| E'/A' | nd | 0.73±0.17 | 0.69±0.05 |
| E/E' | nd | 37.9±3.9 | 38.9±8.4 |
| IVRT (msec) | nd | 27.0±2.9 | 26.6±1.1 |
| VTI (msec) | 22.7±0.9 | 17.0±0.7*** | 19.0±0.8* |
| LVM (mg) | 57.4±3.2 | 90.0±7.1*** | 92.4±5.4*** |
| LVM/BW (mg/g) | 6.5±0.3 | 11.8±0.65**** | 11.2±0.6**** |
| EF (%) | 55.8±4.0 | 28.7±3.3**** | 31.8±3.1*** |
| SV (µl) | 15.9±1.4 | 9.3±1.4** | 10.2±1.4* |
| CO (ml/min) | 7.8±0.8 | 9.4±0.9 | 8.7±0.8 |
| **1 week treatment / 4-week-old** | | | |
| Parameters | WT-Empty | KI-Empty | KI-TTL |
| N number | 12 | 12 | 12 |
| F/M | 7/5 | 12/2 | 7/5 |
| BW (g) | 14.1±0.4 | 12.0±0.4** | 13.1±0.5 |
| IVSd (mm) | 0.49±0.04 | 0.54±0.03 | 0.51±0.02 |
| LVPWd (mm) | 0.50±0.03 | 0.59±0.03 | 0.55±0.03 |
| LVEDD (mm) | 3.67±0.06 | 3.76±0.07 | 3.95±0.08* |
| LVESD (mm) | 2.80±0.08 | 3.26±0.08** | 3.48±0.08**** |
| FAC (%) | 42.5±1.6 | 24.5±1.8**** | 22.8±2.2**** |
| AET (msec) | nd | 25.4±1.3 | 28.2±2.0 |
| MV Decel Time (msec) | nd | 20.6±1.0 | 23.8±1.1 |
| MV E/A | nd | 1.23±0.08 | 1.28±0.09 |
| E'/A' | nd | 0.68±0.15 | 0.69±0.05 |
| E/E' | nd | 42.4±8.9 | 40.4±4.5 |
| IVRT (msec) | nd | 21.0±1.4 | 23.8±1.1 |
| VTI (msec) | 23.1±0.7 | 20.8±1.9 | 22.4±1.3 |
| LVM (mg) | 100.6±6.0 | 112.6±6.3 | 116.9±7.1 |
| LVM/BW (mg/g) | 7.2±0.5 | 9.5±0.6* | 9.0±0.6 |
| EF (%) | 51.9±1.9 | 32.9±2.0**** | 37.0±3.1*** |
| SV (µl) | 27.2±1.6 | 13.6±1.2**** | 19.0±2.8* |
| CO (ml/min) | 13.9±0.8 | 7.5±0.6*** | 9.8±1.4* |

| 2 weeks treatment / 5-week-old | | | |
|---|---|---|---|
| Parameters | WT-Empty | KI-Empty | KI-TTL |
| N number | 12 | 12 | 12 |
| F/M | 7/5 | 12/2 | 7/5 |
| BW (g) | 18.4±0.6 | 16.3±0.4** | 17.7±0.3 |
| IVSd (mm) | 0.46±0.03 | 0.57±0.03* | 0.60±0.03** |
| LVPWd (mm) | 0.45±0.02 | 0.60±0.03** | 0.61±0.03** |
| LVEDD (mm) | 3.61±0.08 | 4.14±0.06**** | 4.40±0.04**** # |
| LVESD (mm) | 2.52±0.11 | 3.56±0.07**** | 3.66±0.11**** |
| FAC (%) | 51.7±2.2 | 26.0±1.8**** | 28.9±2.0**** |
| AET (msec) | 35.9±0.9 | 25.8±1.3* | 27.9±1.6* |
| MV Decel Time (msec) | 17.7±1.1 | 18.2±1.2 | 20.4±1.6** |
| MV E/A | 1.14±0.04 | 1.15±0.04 | 1.17±0.05 |
| E'/A' | 0.74±0.14 | 0.63±0.05 | 0.65±0.06 |
| E/E' | 29.4±3.3 | 48:3±5:9 | 39:7±2:7 |
| IVRT (msec) | 13.7±1.1 | 20.2±1.4 | 19.7±1.3 |
| VTI (msec) | 25.1±0.7 | 20.9±0.8** | 24.8±1.5# |
| LVM (mg) | 103.1±8.2 | 151.5±11.2** | 166.5±7.0**** |
| LVM/BW (mg/g) | 5.7±0.5 | 9.3±0.7**** | 9.4±0.3**** |
| EF (%) | 60.5±2.2 | 36.7±1.8**** | 36.3±2.5**** |
| SV (µl) | 33.2±1.6 | 18.1±1.9**** | 18.2±1.4**** |
| CO (ml/min) | 18.1±0.8 | 9.8±1.1**** | 9.5±0.9**** |
| **4 weeks treatment / 7-week-old** | | | |
| Parameters | WT-Empty | KI-Empty | KI-TTL |
| N number | 12 | 12 | 11 |
| F/M | 7/5 | 12/2 | 7/5 |
| BW (g) | 20.6±0.9 | 17.8±0.5* | 20.0±0.5 |
| IVSd (mm) | 0.53±0.03 | 0.62±0.04 | 0.70±0.04** |
| LVPWd (mm) | 0.52±0.03 | 0.62±0.04 | 0.65±0.04 |
| LVEDD (mm) | 3.92±0.07 | 4.26±0.07** | 4.50±0.07**** |
| LVESD (mm) | 3.04±0.09 | 3.71±0.07**** | 3.85±0.09**** |
| FAC (%) | 41.6±2.3 | 25.6±2.4**** | 27.1±1.6*** |
| AET (msec) | 43.0±1.5 | 27.8±1.1**** | 32±2.0**** |
| MV Decel Time (msec) | 15.4±1.2 | 21.0±1.8 | 25.7±2.8 |
| MV E/A | 1.20±0.05 | 1.16±0.03 | 1.19±0.07 |
| E'/A' | 0.81±0.14 | 0.54±0.02 | 0.61±0.03 |
| E/E' | 32.7±2.9 | 45.3±2:4* | 42.6±3.5 |
| IVRT (msec) | 15.5±1.2 | 22.8±0.9**** | 21.7±1.0*** |
| VTI (msec) | 22.9±0.9 | 20.8±0.9 | 22.7±1.7 |
| LVM (mg) | 128.2±6.4 | 160.2±11.7 | 190.3±13.9** |
| LVM/BW (mg/g) | 6.3±0.3 | 8.9±0.5** | 9.6±0.7*** |
| EF (%) | 50.7±3.7 | 32.7±1.9*** | 39.2±2.0* |
| SV (µl) | 36.4±2.5 | 15.1±1.7**** | 22.8±2.0*** |
| CO (ml/min) | 18.8±1.3 | 8.8±1.1**** | 11.3±1.2*** |
| **8 weeks treatment / 11-week-old** | | | |

| Parameters | WT-Empty | KI-Empty | KI-TTL |
|---|---|---|---|
| N number | 12 | 11 | 11 |
| F/M | 7/5 | 12/2 | 7/5 |
| BW (g) | 23.6±1.1 | 21.1±0.5 | 23.0±0.7 |
| IVSd (mm) | 0.58±0.02 | 0.71±0.05 | 0.76±0.05** |
| LVPWd (mm) | 0.38±0.02 | 0.46±0.03** | 0.50±0.02 |
| LVEDD (mm) | 4.1±0.07 | 4.4±0.09 | 4.711**** # |
| LVESD (mm) | 3.21±0.08 | 3.88±0.07**** | 4.11±0.09**** |
| FAC (%) | 44.2±1.8 | 24.41±1.8**** | 23.9±2.0**** |
| AET (msec) | 44.8±2.2 | 31.1±1.2**** | 29.5±1.6**** |
| MV Decel Time (msec) | 19.7±1.6 | 20.3±1.0 | 23.1±1.1 |
| MV E/A | 1.20±0.06 | 1.13±0.06 | 1.24±0.03 |
| E'/A' | 0.65±0.08 | 0.66±0.10 | 0.56±0.06 |
| E/E' | 35.8±2.7 | 42.2±3.3 | 48.1±3.6* |
| IVRT | 16.3±0.4 | 22.8±1.0**** | 23.1±0.7**** |
| VTI (msec) | 24.1±1.2 | 23.0±1.2 | 27.8±1.9 |
| LVM (mg) | 133.2±6.6 | 220.2±9.8**** | 242.4±14.1**** |
| LVM/BW (mg/g) | 5.7±0.3 | 10.5±0.5**** | 10.5±0.5**** |
| EF (%) | 54.5±1.1 | 28.3±2.4**** | 38.0±1.3**** ## |
| SV (µl) | 39.7±1.7 | 17.0±2.3**** | 25.4±2.0**** # |
| CO (ml/min) | 20.2±1.0 | 8.6±1.2**** | 12.4±1.1**** |
| **12 weeks treatment - 15-week-old** | | | |
| | WT-Empty | KI-Empty | KI-TTL |
| N number | 12 | 11 | 11 |
| F/M | 7/5 | 12/2 | 7/5 |
| BW (g) | 25.6±1.0 | 22.6±0.6* | 24.0±0.7 |
| IVSd (mm) | 0.64±0.03 | 0.69±0.05 | 0.63±0.04 |
| LVPWd (mm) | 0.63±0.03 | 0.74±0.05 | 0.67±0.05 |
| LVEDD (mm) | 3.93±0.07 | 4.33±0.10* | 4.85±0.22**** # |
| LVESD (mm) | 2.76±0.10 | 3.82±0.06**** | 4.16±0.12**** |
| FAC (%) | 44.3±2.9 | 26.4±2.2**** | 24.4±2.3**** |
| AET (msec) | 37.9±1.5 | 26.8±1.2**** | 29.8±1.5*** |
| MV Decel Time (msec) | 20.5±1.8 | 18.0±1.5 | 17.6±1.6 |
| MV E/A | 1.10±0.05 | 1.10±0.08 | 1.29±0.11 |
| E'/A' | 0.79±0.12 | 0.75±0.15 | 0.76±0.17 |
| E/E' | 32.0±3.0 | 33.4±5.6 | 45.9±3.6 |
| IVRT (msec) | 12.3±0.7 | 21.5±1.3**** | 23.3±0.6**** |
| VTI (msec) | 24.0±1.0 | 19.8±1.4 | 22.4±2.1 |
| LVM (mg) | 134.9±4.3 | 231.7±8.9*** | 247.4±12.9*** |
| LVM/BW (mg/g) | 5.3±0.2 | 10.3±0.4**** | 10.3±0.3**** |
| EF (%) | 60.7±2.8 | 28.0±2.0**** | 40.3±3.5**** # |
| SV (µl) | 40.6±2.7 | 16.2±1.6*** | 28.7±2.7** ## |
| CO (ml/min) | 22.8±1.5 | 8.8±0.9**** | 14.5±1.4*** # |

Data are expressed as mean±SEM. *P<0.01, **P<0.01, ***P<0.001, ***P<0.0001 vs. WT-Empty, and #P<0.01, ##P<0.01 vs. KI-Empty, one-way ANOVA, followed by Tukey's multiple comparisons test. Abbreviations: **AET,** aortic ejection time; **BW,** body weight; **CO,** cardiac output; **FAC,** fractional area change; **E'/A',** peak velocity blood flow from LV relaxation in early diastole (E´ wave) to peak velocity flow in late diastole caused by atrial contraction (A' wave) obtained by tissue Doppler of the mitral annulus; **E/E',** calculated ratio of peak velocity in early diastole; **EF,** ejection fraction; **F/M,** number of females/males; **IVRT,** isovolumic relaxation time; **IVSd,** interventricular septum thickness in diastole; **LVM,** left ventricular mass; **LVM/BW,** left ventricular mass to body weight ratio; **N,** number of mice; **nd,** not determined; **LVEDD,** left ventricular end-diastolic diameter; **LVESD,** left ventricular end-systolic diameter; **LVPWd,** left ventricular posterior wall thickness in diastole; **MV E/A,** peak velocity blood flow from LV relaxation in early diastole (E wave) to peak velocity flow in late diastole caused by atrial contraction (A wave) obtained by Tissue Doppler through the mitral valve; **MV Decel Time,** mitral valve deceleration time; **SV,** stroke volume; **VTI,** velocity time interval.

**Table S3. Hemodynamic parameters in *Mybpc3*-targeted knock-in (KI) and wild-type (WT) mice treated with AAV9-TTL and/or AAV9-Emtpy.**

| **12 weeks treatment / 15-wk-old** | | | |
|---|---|---|---|
| Parameters | WT-Empty | KI-Empty | KI-TTL |
| N number | 9 | 9 | 11 |
| F/M | 5/4 | 7/2 | 7/5 |
| BW (g) | 24.7±1.2 | 22.5±0.5 | 23.7±0.8 |
| **Global function** | | | |
| Heart rate (bpm) | 581±12 | 564±10 | 572±11 |
| Cardiac output (ml/min) | 16.37±1.9 | 10.71±1.09* | 15.29±1.52# |
| Stroke volume (µl) | 28.3±3.3 | 18.9±1.9* | 26.9±2.7# |
| Stroke work (mmHg*µl) | 2107±261 | 1250±194* | 1986±249# |
| **Systolic function** | | | |
| Ejection fraction (%) | 80±6 | 49±4*** | 51±5** |
| LVESP (Pes) (mmHg) | 68±4 | 78±2 | 81±4 |
| $dP/dt_{max}$ (mmHg/s) | 10778±908 | 9351±778 | 11422±829 |
| LVESV (µl) | 9.7±4 | 25.7±2.6 | 35.5±6.1 |
| $dV/dt_{max}$ (µl/s) | 1922±222 | 1564±210 | 1867±230 |
| PRSW (mmHg) | 87±7.5 | 65.2±8:2 | 79.5±4:9 |
| **Diastolic function** | | | |
| LVEDP (Ped) (mmHg) | 2.37±0.42 | 3.64±0.54 | 2.95±0.52 |
| $dP/dt_{min}$ (mmHg/s) | -8448±652 | -4738±216**** | -5372±322*** |
| LVEDV (µl) | 32.7±5.5 | 36.8±3.1 | 54.8±6.3*# |
| $dV/dt_{min}$ (µl/s) | -2208±218 | -2158±260 | -3024±313 |
| Tau (ms) | 4.62±0.18 | 7.28±0.31**** | 6.48±0.32*** |
| EDPVR (mmHg/µl) | 0.29±0.07 | 0.34±0.04 | 0.18±0.04# |

Data are expressed as mean ± SEM; *P<0.05, **P<0.01, ***P<0.001, ****P<0.0001, Student's t-test vs. WT and #P<0.05, ##P<0.01, ###P<0.001, Student's t-test vs. KI-Empty. Abbreviations: **BW,** body weight; **$dP/dt_{max}$,** maximal rate of left ventricular pressure development in systole; **$dP/dt_{min}$,** maximal rate of left ventricular pressure development in diastole; **$dV/dt_{max}$,** point of maximum volume increase; **$dV/dt_{min}$,** point of maximum volume decrease; **EDPVR,** end-diastolic pressure-volume relation; **F/M,** number of females/males; **LVEDV,** left ventricular end-diastolic volume; **LVESP,** left ventricular end-systolic pressure; **N,** number of mice; **PRSW,** preload recruitable stroke work; **Tau,** time constant of active relaxation.
